# GDF15 and ACE2 stratify COVID19 patients according to severity while ACE2 mutations increase infection susceptibility

**DOI:** 10.1101/2022.05.06.490907

**Authors:** Margalida Torrens-Mas, Catalina M Perelló-Reus, Neus Trias-Ferrer, Lesly Ibargüen-González, Catalina Crespí, Aina Maria Galmes-Panades, Cayetano Navas-Enamorado, Andres Sanchez-Polo, Javier Piérola-Lopetegui, Luis Masmiquel, Lorenzo Socias Crespi, Carles Barcelo, Marta Gonzalez-Freire

**Affiliations:** Translational Research in Aging and Longevity Group (TRIAL group), Health Research Institute of the Balearic Islands (IdISBa), Palma de Mallorca, Spain; Translational Pancreatic Cancer Oncogenesis Group, Health Research Institute of the Balearic Islands (IdISBa), Palma de Mallorca, Spain; Cell Culture and Flow Cytometry Facility, Health Research Institute of the Balearic Islands (IdISBa), Palma de Mallorca, Spain; Physical Activity and Sport Sciences Research Group (GICAFE). Institute for Educational Research and Innovation (IRIE). University of the Balearic Islands, Palma de Mallorca, Spain; Microscopy Facility, Health Research Institute of the Balearic Islands (IdISBa), Palma de Mallorca, Spain; Vascular and Metabolic Pathologies Group, Health Research Institute of the Balearic Islands (IdISBa), Palma de Mallorca, Spain; Intensive Care Unit, Son Llatzer University Hospital, Balearic Islands, Palma de Mallorca, Spain

## Abstract

Coronavirus disease 19 (COVID-19) is a persistent global pandemic with a very heterogeneous disease presentation ranging from a mild disease to dismal prognosis. Early detection of sensitivity and severity of COVID-19 is essential for the development of new treatments. In the present study, we measured the levels of circulating growth differentiation factor 15 (GDF15) and angiotensin-converting enzyme 2 (ACE2) in plasma of severity-stratified COVID-19 patients and healthy control patients and characterized the in vitro effects and cohort frequency of *ACE2* SNPs. Our results show that while circulating GDF15 and ACE2 stratify COVID-19 patients according to disease severity, *ACE2* missense SNPs constitute a risk factor linked to infection susceptibility.

## 1. Introduction

In December 2019, several cases of pneumonia emerged in Wuhan, China, which were caused by a novel coronavirus initially named 2019-nCoV (1). Due to its phylogenetic proximity to SARS-CoV, this new coronavirus was renamed as ‘SARS-CoV-2’ by the International Committee on Taxonomy of Viruses (2). This virus rapidly spread around the world due to its high transmissibility and the presence of asymptomatic subjects, which led to the declaration of a global pandemic by the World Health Organization. COVID-19 shows a very heterogeneous disease presentation, and clinical symptoms may differ with age and sex. Disease severity has been associated with older age, male sex, and comorbidities such as hypertension, diabetes, obesity, cardiovascular disease, chronic obstructive pulmonary disease, and lung, liver, and kidney disease (3–5). Severe or fatal cases of COVID-19 usually present with increased levels of pro-inflammatory cytokines and chemokines, known as cytokine storm (6), low albumin levels (7), as well as decreased lymphocyte counts and platelets, elevated levels of C-Reactive Protein (CRP), procalcitonin, D-dimer, lactate dehydrogenase, and ferritin, among others (8–10). Similarly, recent studies report that GDF15 levels, a member of the transforming growth factor-β (TGF-β) superfamily, are increased in COVID-19 patients who require hospitalization, and its levels are associated with viremia, hypoxemia, and worse clinical outcome (11–13). It is known that GDF15 levels are upregulated by inflammation, cancer, cardiovascular and metabolic diseases (14, 15). Also, circulating GDF15 levels show a high correlation with age and are primarily expressed under conditions of inflammation and oxidative stress (16). On the other hand, a predisposing genetic background may contribute to the wide clinical variability of COVID-19. To date, genetic markers of susceptibility to COVID-19 have not yet been identified.

The most well-known and evaluated mechanism of SARS-CoV-2 infection is the binding and uptake of viral particle through the ACE2 receptor, a type 1 integral membrane glycoprotein. The spike protein of the SARS-CoV-2 virus mediates its entry into the host cell through its interaction with ACE2. Transmembrane protease serine 2 (TMPRSS2) also participates in the cleavage of the spike protein, facilitating the entry. Both ACE2 and TMPRSS2 are highly expressed in different tissues and cell types, including the type II alveolar epithelial cells (17, 18). It has been observed that ACE2 expression in this type of cells increases with age, which could contribute to the severity of COVID-19 symptoms among elderly patients (19, 20). For instance, plasma levels of ACE2 have been found elevated in kidney disease and cardiovascular disease (21). Recent studies show that ACE2 plasma levels are increased in COVID-19 patients compared to healthy subjects (22, 23).

The emergence of SARS-CoV-2 variants-of-concern from last semester of 2021, accounted for vast majority of reported worldwide COVID-19 positivity. Such variants represent a public health challenge during the COVID-19 pandemic due to their ability to increase viral transmission and disease severity. Several mechanisms might account for increased variant transmissibility, especially mutations involved in enhanced spike protein binding affinity for the *ACE2* receptor (24). Therefore, coding variants within *ACE2* could have the potential to alter SARSCoV-2 binding and possible infection responses in different individuals based on host genetics (25–28).

Based in all the above, the aims of this study were: i) to analyze whether GDF15 and ACE2 levels correlate with a worse COVID-19 prognosis as well as with other inflammatory and cellular markers of damage and senescence, and ii) to determine if different SNPs variants within *ACE2* were associated to disease severity.

For this, circulating levels of both proteins, as well as RNA and DNA, were analyzed from plasma and buffy coats samples obtained from COVID-19 patients and in a healthy control group matched by age and sex.

## 2. Results

### 2.1 Demographic and clinical characteristics

Demographic data, comorbidities, and inflammation markers of the individuals included in this study are shown in Table 1. A total of 72 subjects were included in the study, 46% males, with an average age of 69 years (rank 28-88 years). SARS-CoV-2 infection was diagnosed by a real-time RT-PCR. There were no differences in age or sex among the three groups (p>0.05). All the ICU patients died. The most frequent comorbidities were hypertension (51%), type 2 diabetes mellitus (38%), smokers (38%), or others (59%). Inflammatory markers such as ferritin, CRP, and D-dimer were significantly higher in the ICU group that died, compared to the non-ICU and control group.

**Table 1.** Baseline characteristics of the sample population.

|  | Total (n=72) | ICU (n=21) | non-ICU<br>(n=26) | Control group<br>(n=25) | p-value |
| --- | --- | --- | --- | --- | --- |
| **Age (years)** | 69.51(10.31)<br>[62-72] | 71.2 (13.87)<br>[65-78] | 71.84 (7.38)<br>[69-75] | 65.84 (8.78)<br>[62-69] | 0.081 |
| **Males (%)** | 33 (46%) | 10 (48%) | 13 (50%) | 10 (40%) | 0.759 |
| **Mortality (%)** | 21 (29.16%) | 21 (100%) | 0 (0%) | 0(0%) | <0.001* |
| **Comorbidities** |  |  |  |  |  |
| **Hypertension** | 29/57<br>(51%) | 6/11 (54%) | 15/26 (58%) | 8/20 (40%) | 0.475 |
| **Type 2 diabetes** | 22/57<br>(38%) | 6/11 (54%) | 11/26 (42%) | 5/20 (25%) | 0.236 |
| **Obesity** | 7/57<br>(12%) | 2/11 (18%) | 3/26 (42%) | 2/20 (10%) | 0.792 |
| **COPD** | 3/57 (5) | 3/11 (27%) | 0/26 (0%) | 0/20 (0%) | 0.001* |
| **Dyslipidemia** | 18/57<br>(31%) | 4/11 (36%) | 14/26 (54%) | 0/20 (0%) | <0.001* |
| **Other** | 33/56<br>(59%) | 7/11 (64%) | 14/26 (54%) | 12/19 (63%) | 0.772 |
| **Smokers** | 22/57<br>(38%) | 5/11 (45%) | 17/26 (65%) | 0/20 (0%) | <0.001* |
| **Inflammation markers** |  |  |  |  |  |
| **Ferritin (ng/mL)** | 651.64<br>(1068.24)<br>[327-976] | 1332.79<br>(1350.07)<br>[682-1984] | 133.93<br>(176.17) [61-207] | - | <0.001* |
| **CRP (mg/dL)** | 39.93<br>(78.40) [16-64] | 89.65<br>(98.31) [42-137] | 0.56 (0.82)<br>[0.22-0.91] | - | <0.001* |
| **D-Dimer (ng/mL)** | 907.98<br>(2057.98)<br>[275-1541] | 1761.25<br>(2810.61)<br>[446-3077] | 166.00<br>(160.64)<br>[97-235] | - | <0.001* |
| **GDF15 (pg/mL)** | 2079.88<br>(1594.51)<br>[6606-9838] | 3846.84<br>(1473.42)<br>[3176-4518] | 1620.29<br>(972.05)<br>[1228-2013] | 1073.60 (844.13)<br>[725-1422] | <0.001* |
| **ACE2 (pg/mL)** | 8221.84<br>(6826.93)<br>[6606-9838] | 10107.36<br>(5337.96)<br>[7678-12537] | 7172.9<br>(9243.67)<br>[3439-10906] | 7708.50 (4407.88)<br>[5847-9570] | 0.011* |

Data are presented as mean, standard deviation (), rank [], or percentage (%). \*Statistically significant difference. Some patients' inflammation markers or comorbidities were not available. In this case, total number of patients with available data is shown in each row.

### 2.2 Circulating levels of GDF15 and ACE2

Plasma levels of both GDF15 and ACE2 were measured using ELISA (Figure 1A-B). ICU patients showed higher levels of GDF15 when compared to non-ICU patients and control group (3846.84 pg/mL vs 1620.29 pg/mL and 1073.6 pg/mL, respectively; *P*<0.001; Figure 1A). However, there was no difference between the non-ICU and the control healthy group (*P*>0.05). ACE2 circulating levels (Figure 1B) were increased in the ICU group compared with the non-ICU and control groups (10107.36 pg/mL vs 7172.9 pg/mL and 7708.5 pg/mL, respectively). ICU patients showed significantly higher levels than the non-ICU patients (*P*=0.02). Interestingly, the ratio between the levels of GDF15 and the levels of ACE2 (Figure 1C) was also significantly higher in the ICU patients compared to the non-ICU and control groups (*P*=0.010 and *P*<0.001, respectively). This ratio was also increased in non-ICU patients compared to the control group (*P*=0.006). Finally, we found a positive correlation, although not statistically significant, between the levels of GDF15 and ACE2 (Figure 2), (r=0.204, *P*=0.087).

**Figure 1.**
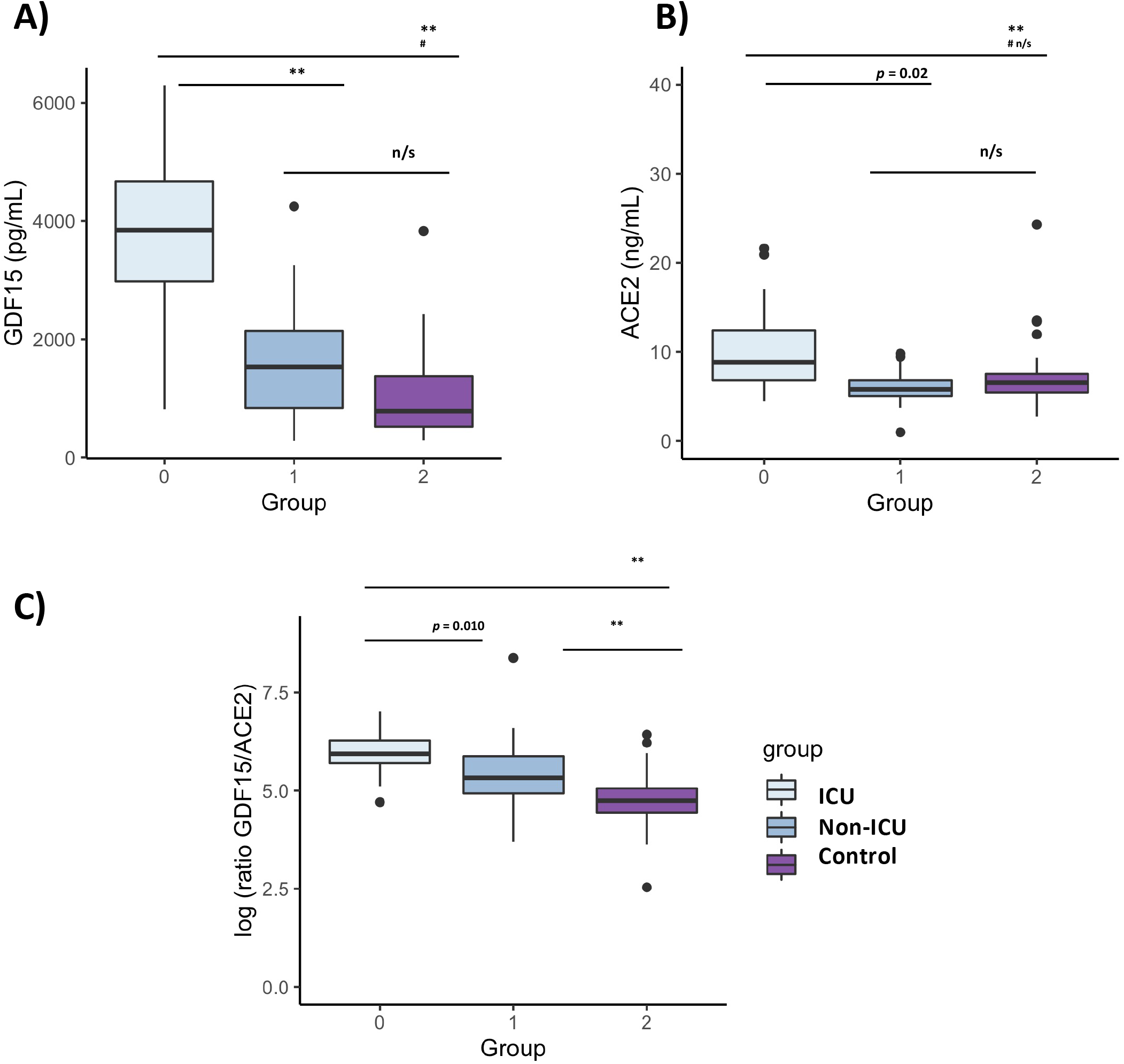
Differences in circulating GDF15 and ACE2 levels in COVID-19 patients and a healthy control group. A) GDF15 levels; B) ACE2 levels ; C) ratio GDf15/ACE2 ; The box plots represent the maximum and minimum levels (whiskers), the upper and lower quartiles, and the median. The length of each box represents the interquartile range. Dots represent outliers. Statistical significance between groups was determined using the ANOVA test. ** p <0.001, # p <0.001 UCI vs Control, n/s (non significant); GDF15 (Growth differentiation factor 15); ACE2 (angiotensin-converting enzyme 2).

**Figure 2.**
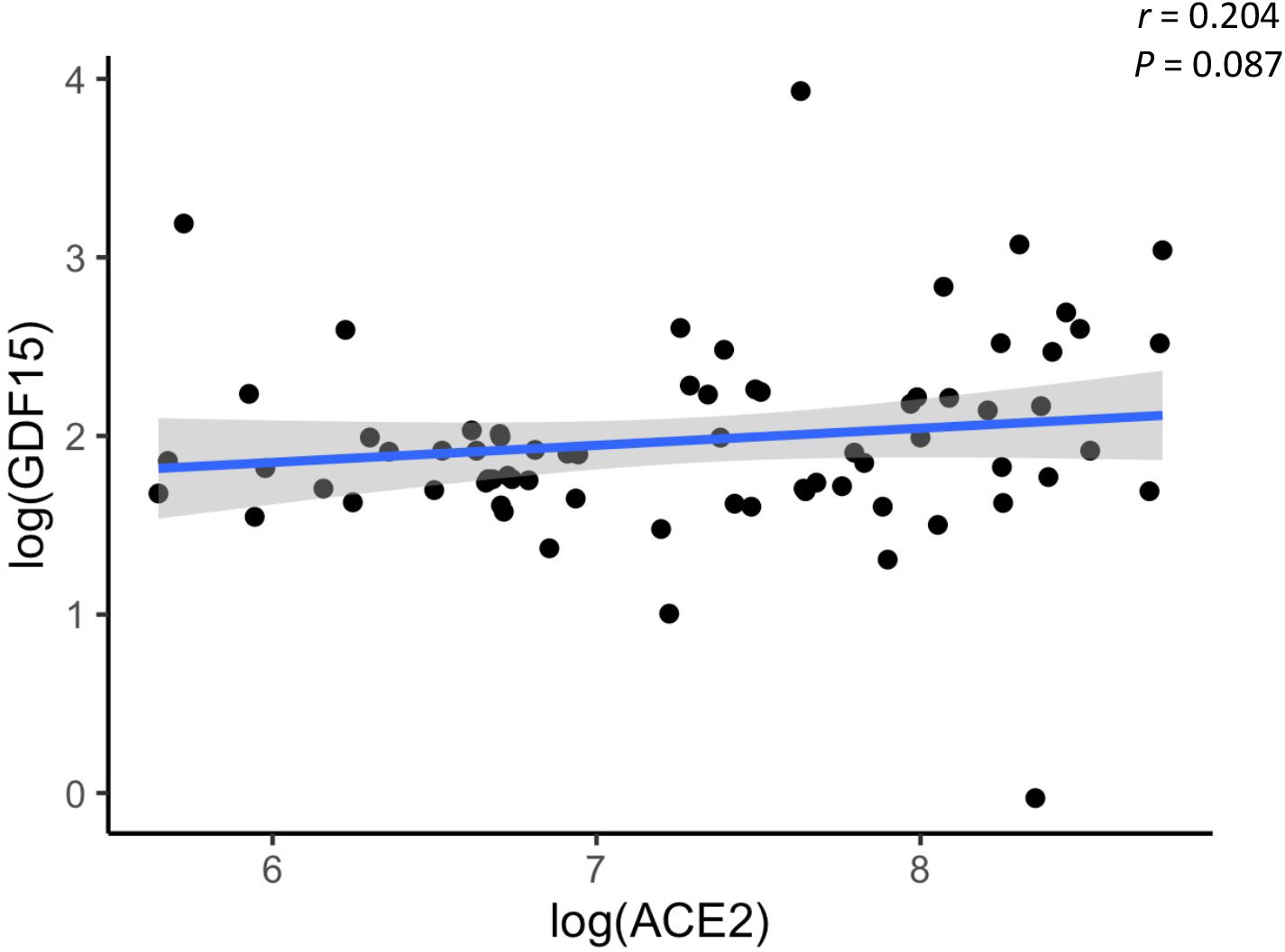
Representative scatterplot showing the association between GDF15 and ACE2 levels. Results of the 72 patients and healthy subjects are shown. Each dot represents an individual value. The solid blue line represents the regression line. The grey shade represents the confidence interval.

### 2.3. Association between the levels of GDF15 and ACE2, with age, sex, and inflammatory markers

As expected, circulating GDF15 was positively associated with age (r=0.482, *P*<0.001, Figure 3A). However, ACE2 levels were not associated with age (r=-0.003, *P*=0.982, Figure 3B). No differences were found in the circulating levels of GDF15 (Figure 4A) and ACE2 (Figure 4B) with sex (*P* >0.05). Since ACE2 is the functional receptor of SARS-CoV-2, we checked whether there were differences in the ACE2 mRNA by RT-qPCR (Figure 5). ICU patients showed higher levels of ACE2 expression compared to non-ICU and control groups (*P*=0.006 and *P*=0.003, respectively). Interestingly, ACE2 expression levels did not correlate with ACE2 protein levels measured by ELISA (data not shown).

**Figure 3.**
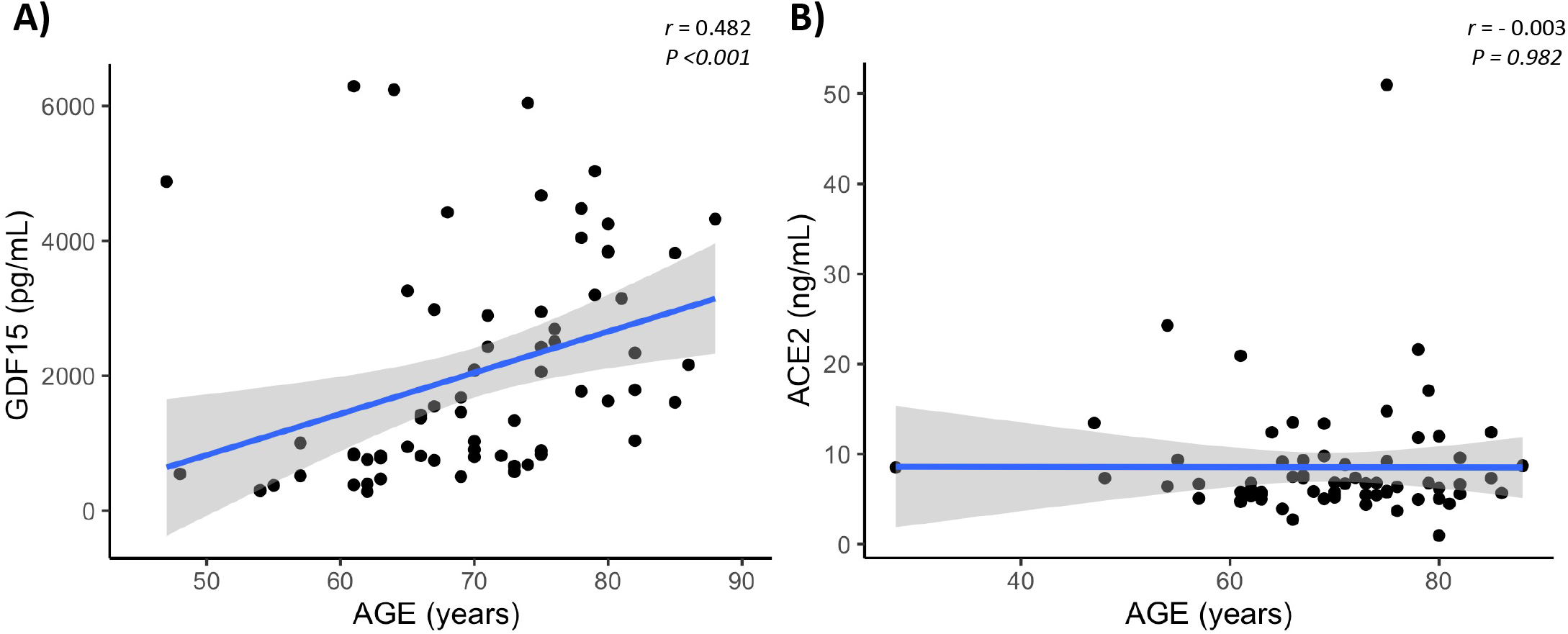
Representative scatterplots showing the association between GDF15 and ACE2 levels with age. GDF15 is positive associated with age (A), while ACE2 is not correlated with age (B). Each dot represents an individual value. The solid blue line represents the regression line. The grey shade represents the confidence interval.

**Figure 4.**
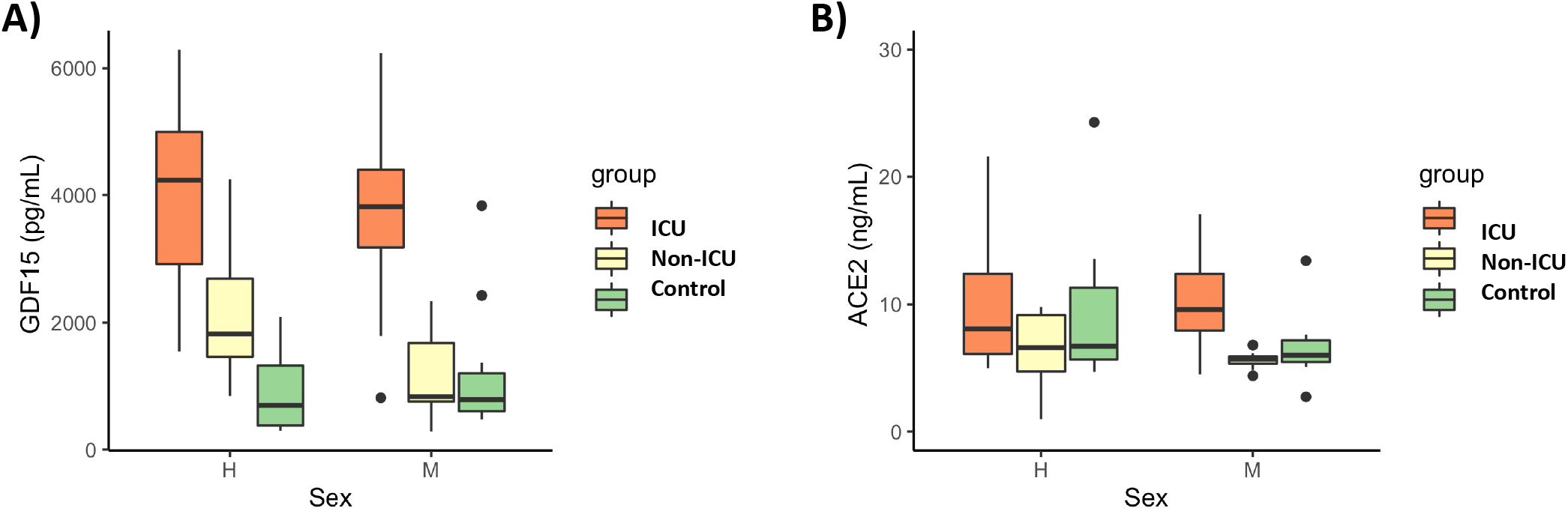
Differences in circulating GDF15 and ACE2 levels by sex in COVID-19 patients and the healthy control group. No differences were found by sex in GDF15 and ACE levels. The box plots represent the maximum and minimum levels (whiskers), the upper and lower quartiles, and the median. The length of each box represents the interquartile range. Dots represent outliers. Statistical significance between groups was determined using the ANOVA test.

**Figure 5.**
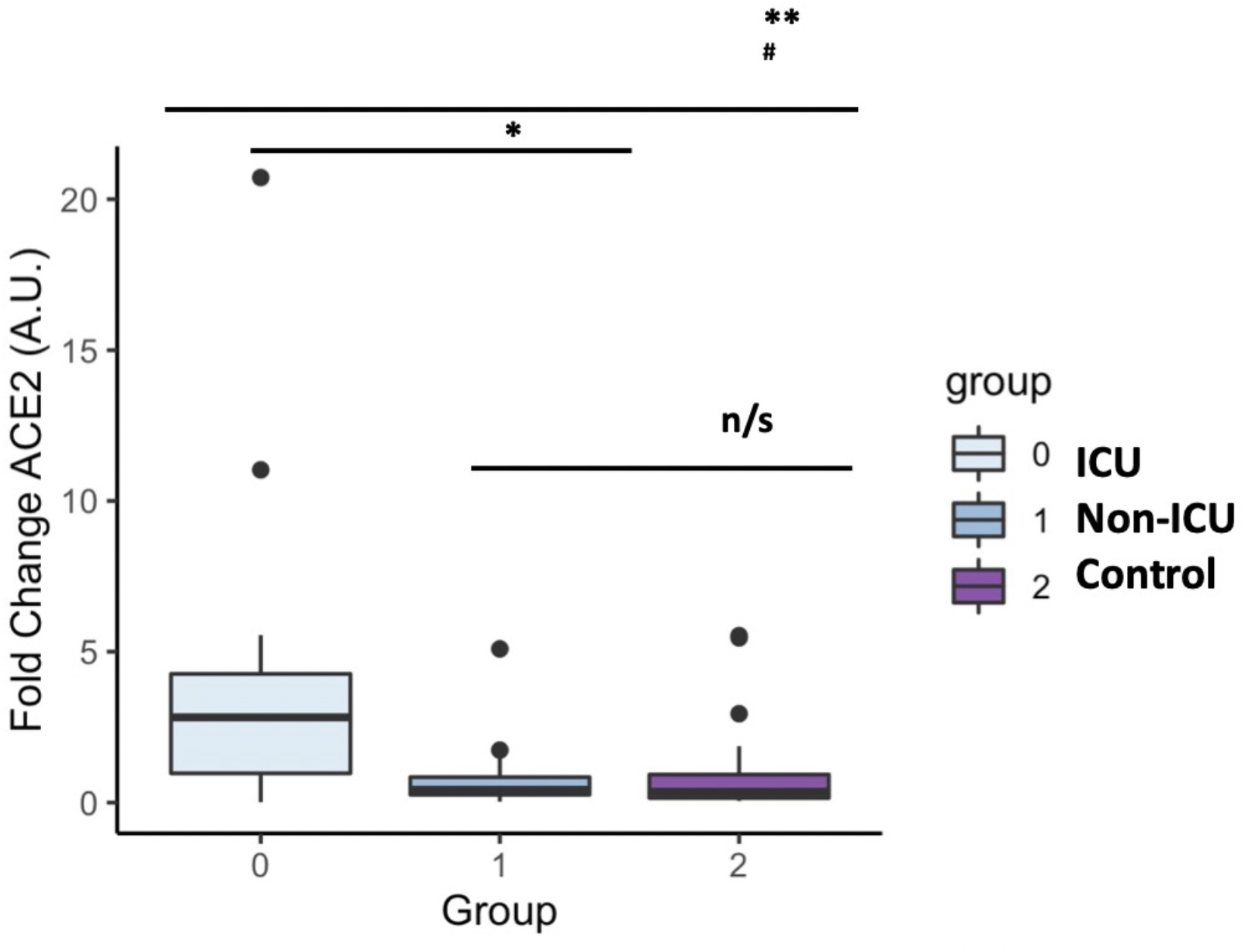
ACE2 mRNA levels in COVID-19 patients and the healthy control group. The box plots represent the maximum and minimum levels (whiskers), the upper and lower quartiles, and the median. The length of each box represents the interquartile range. Dots represent outliers. Statistical significance between groups was determined using the ANOVA test. ** p <0.001, # p <0.001 UCI vs Control, n/s (non significant); ACE2 (angiotensin-converting enzyme 2).

We next analyzed the association between the levels of GDF15 and ACE2 with ferritin, D-dimer and CRP, classical markers of inflammation. Only COVID-19 patients had data on inflammatory markers. As expected, the levels of both GDF15 (Figure 6A) and ACE2 (Figure 6B) were positively correlated with the levels of ferritin, D-dimer, and CRP (GDF15: r=0.561, r=0.688, r=0.497, respectively, *P*<0.001; ACE2: r= 0.386 and *P*=0.009, r=0.350 and *P*=0.021, r=0.381 and *P*=0.012, respectively).

**Figure 6.**
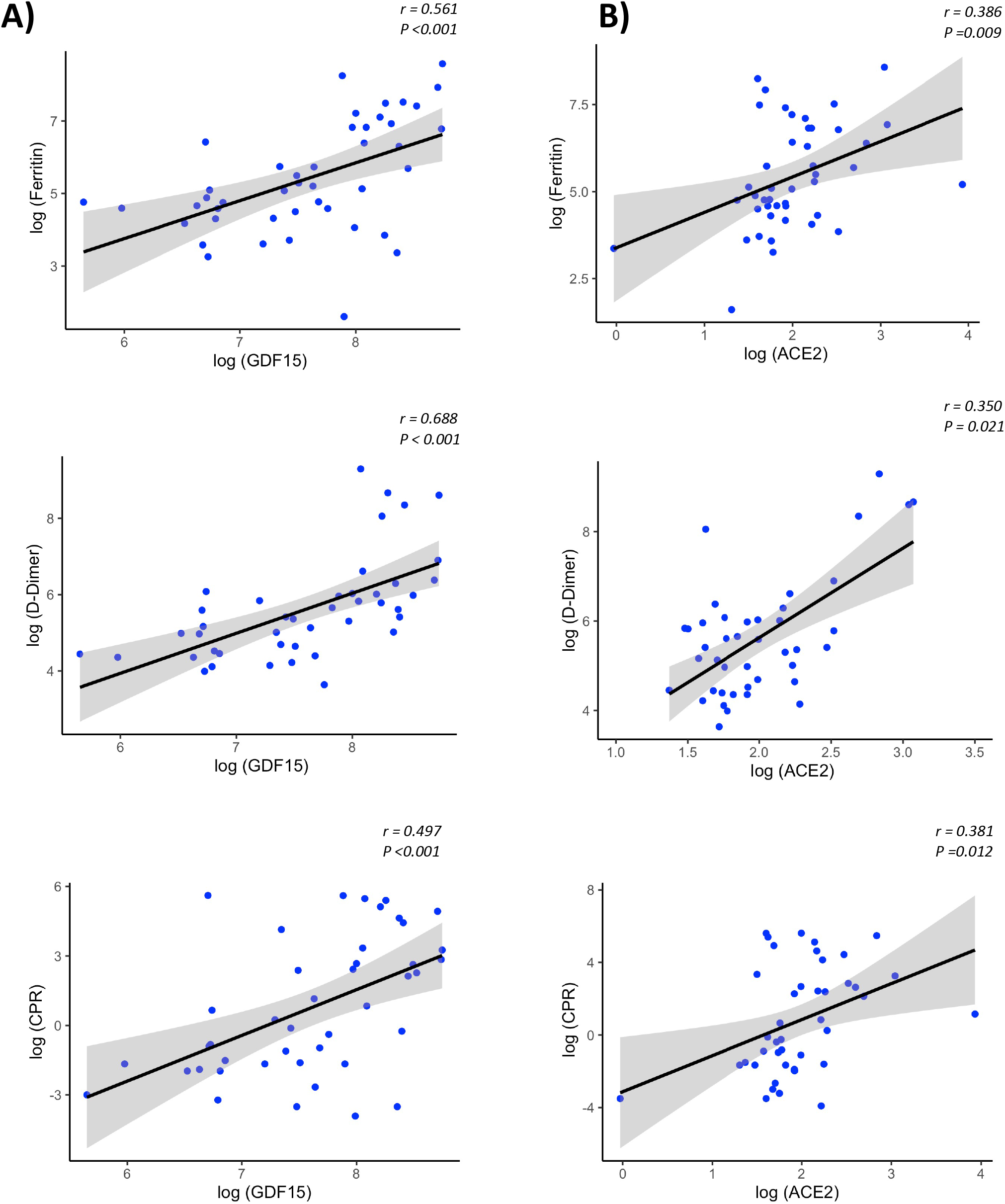
Representative scatterplots (spearman correlations) showing the association between GDF15 and ACE2 levels with classical inflammatory markers. .The solid line represents the regression line. The grey shade represents the confidence interval.

Since hypoxia is one of the main pathophysiological causes of mortality in patients with COVID-19 and the release of damaged mitochondrial DNA (*mtDNA*) is related to the increase of the systemic inflammatory response, we sought to determine the circulating levels of hypoxia inducible factor 1 A (*HIF1A*) mRNA, a transcription factor that normally is upregulated during hypoxic and inflammatory conditions, and circulating levels of *mtDNA*, measured as the ratio between nuclear to mitochondrial DNA. Unexpectedly, we did not find differences neither in the expression levels of *mtDNA* or *HIF1A* (Figure 7A and 7B respectively).

**Figure 7.**
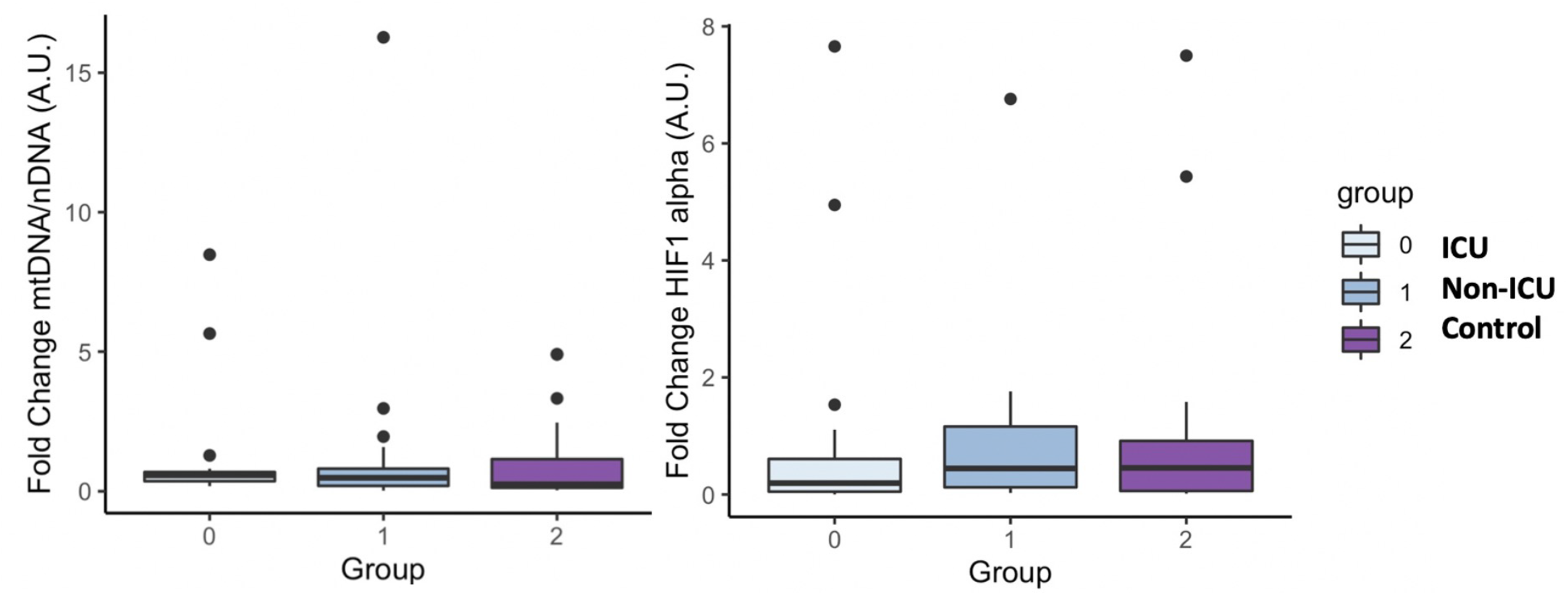
mtDNA and HIF1 alpha mRNA levels in COVID-19 patients and the healthy control group. The box plots represent the maximum and minimum levels (whiskers), the upper and lower quartiles, and the median. The length of each box represents the interquartile range. Dots represent outliers. Statistical significance between groups was determined using the ANOVA test. No difference were found in mtDNA and HIF1 alpha mRNA levels among groups.

We performed principal component analysis (PCA) with all the markers above plus GDF15, ACE2, age, and sex (Figure 8). These analyses showed a clear distinction between ICU, non-ICU patients and the control group, revealing a clear effect of SARS-CoV-2 infected patients compared to healthy subjects.

**Figure 8.**
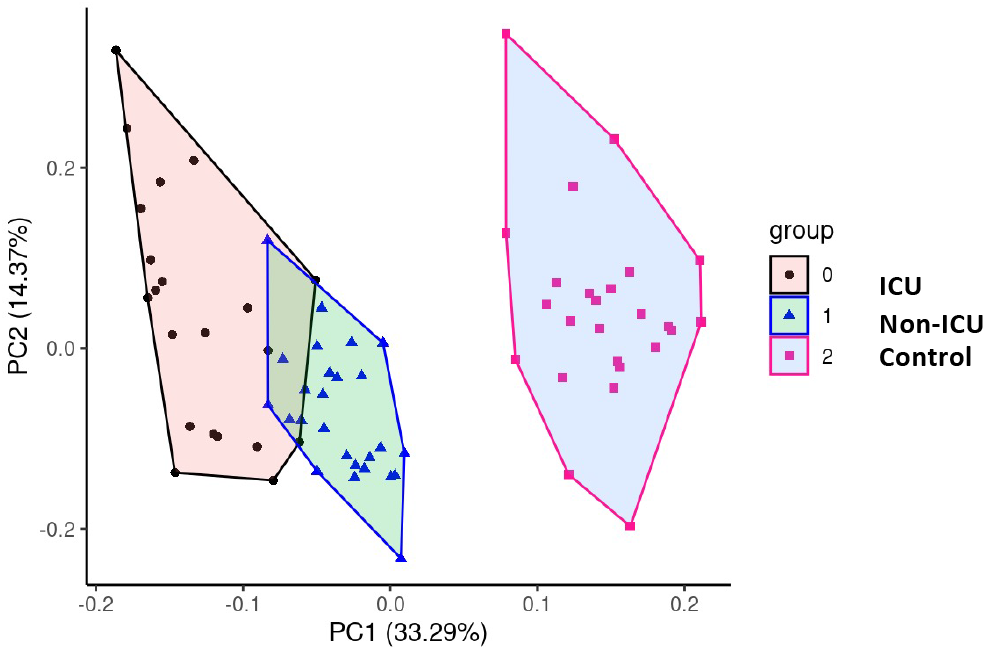
Principal Component Analysis (PCA) To test potential associations between the studied parameters, a PCA was computed, showing a clear distinction between COVID-19 patients and healthy subjects.

We had complete clinical data from blood routine analysis (Supplemental Table 3) as well as buffy coats from the non-UCI group, we decided to test if the levels of GDF15 and ACE2 were correlated with those parameters. We also extracted RNA and DNA from buffy coats and measured known markers of senescence and oxidative damage, such as *p16, p21*, telomere length, *mtDNA/nDNA* ratio and *mtDNA* oxidation. In the Supplemental Table 4 we show the correlation coefficient between all the variables described above, GDF15 and ACE2. Interestingly, both circulating GDF15 and ACE2 were positively correlated with glucose, urea, creatinine, and N-terminal (NT)-pro hormone BNP (NT-proBNP) (*P*<0.01). Circulating GDF15 was negatively correlated to the platelets levels (r= −0.557, *P* <0.05) and the levels of *mtDNA* oxidation (r= −0.495, *P* <0.05). With regards to mRNA expression data, the expression levels of *ACE2* in plasma were also correlated with the neutrophil and the ratio lymphocyte to neutrophils (r= 0.478, p <0.05; r= 0.629, *P*<0.01 respectively), while the expression levels of ACE2 in buffy coats were only associated with the eosinophils levels (r= 0.542, *P* <0.05). Finally, the protein and mRNA expression levels of *ACE2* in plasma were highly positive correlated with the expression of *HIF1A* in buffy coats (r= 0.637, *P* <0.05), while the levels of *ACE2* in buffy coats were correlated with the expression of *HIF1A* in plasma (r= 0.704, *P* <0.01).

### 2.4. Differential SARS-CoV-2 infection dependent on COVID-19 variant-of-concern and ACE2 expression

Since ACE2 circulating levels were increased in the ICU group compared to the non-ICU and control group, both at mRNA and protein levels (Figures 5 and 1B), we then tested the effect of full length *ACE2* expression on the infection capacity of SARS-CoV-2. To this end, we developed an infection model based on SARS-CoV-2 pseudotyped lentivirus (pseudovirus) expressing Spike protein and encapsulating a mCherry reporter. We tested infection with this approach in human alveolar lung A549 cells, as a COVID-19 human infection model to recapitulate airway infection (Figure 9A). We found untransfected A549 to be non-permissive to SARS-CoV-2 pseudoinfection (Figure 9B) due to the low expression of the endogenous viral receptor *ACE2* (Supplemental Figure 1). This eliminates the interference of endogenous ACE2 when overexpressing *ACE2* variants and facilitates the interpretation of results when performing pseudovirus entry studies. Exogenous expression of GFP-*ACE2* enabled SARS-CoV-2 pseudovirus to effectively infect the cells with mCherry reporter as determined by Fluorescent Activated Cell Sorter (FACS) (Figure 9B).

**Figure 9.**
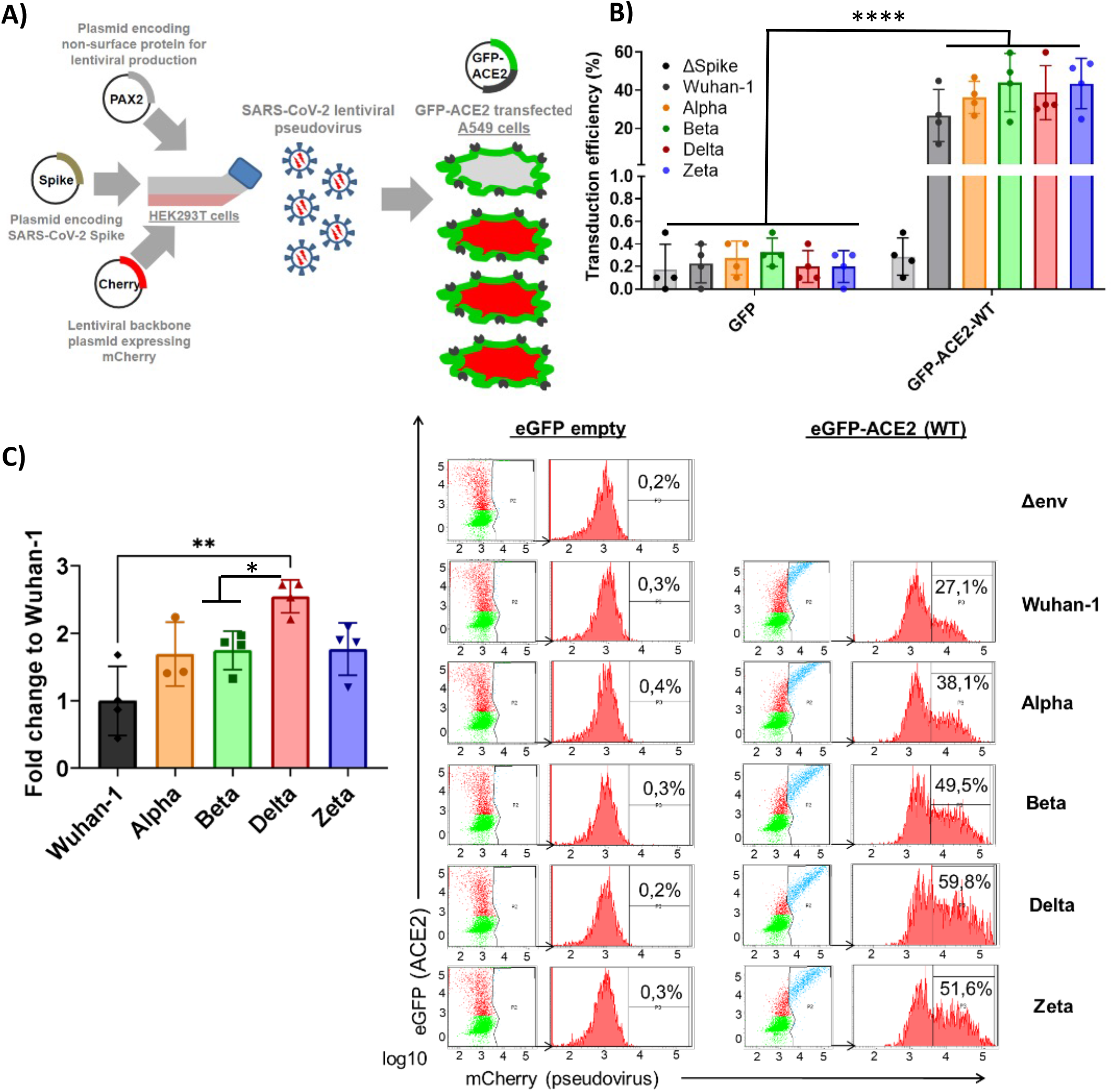
Differential SARS-CoV-2 infection is dependent on COVID19 variant-of-concern and ACE2 expression. A) SARS-CoV-2 cell entry assay strategy: Lentiviral-based replication-defective pseudovirus were generated in HEK293T cells from lentiviral parental genes, SARS-CoV-2 Spike and encapsulating a mCherry reporter. Since the entry steps of the SARS-CoV-2 pseudovirions are governed by the coronavirus Spike protein at their surface, they enter cells in a similar fashion to native counterparts. A549 airway cells were transfected with exogenous GFP-hACE2 enabling SARS-CoV-2 pseudovirus to effectively infect the cells with mCherry reporter. Double-positive GFP/mCherry cells were quantified by flow cytometry to assess viral infection capacity. B,C) Following the strategy described in A, A549 cells expressing GFP-hACE2 were assayed for cell entry by SARS-CoV-2 pseudovirus expressing either empty vector (Δ Spike) or Spike protein corresponding to origin variant (Wuhan-1) or variants-of-concern Alpha, Beta, Delta or Zeta. A representative flow cytometry experiment is shown. Bars demonstrate mean and Standard Error of Mean while each data point represents a unique experiment; ****P < 0.0001; **P < 0.01; *P < 0.1 by t test.

Finally, in order to test whether the main variants of concern (VOC) circulating the last semester of 2021 (B.1.1.7 (Alpha), B.1.351 (Beta), P.2 (Zeta) and B.1.617.2 (Delta)) infected ACE2-expressing cells with differential efficiency, we expressed the different full-length Spike protein corresponding to the VOCs (Wuhan-1, Alpha, Beta, Delta and Zeta) in the lentiviral pseudovirus model described above and infected the A549 transiently expressing full length *ACE2* described above.

We found significant increased infectivity in the Delta variant as compared to the origin variant Wuhan-1 (*P<*0.016). Alpha, Beta and Zeta showed a modest non-significant increased infectivity as compared to Wuhan-1 (Figure 9C).

### 2.5. *ACE2* polymorphisms K26R, P389 and N720D promote SARS-CoV-2 infection in all VOCs

Given that host genetics and ethnicity predisposition to COVID-19 is now recognized to be relevant to COVID-19 susceptibility and severity, we reasoned that it is very likely that there exists *ACE2* variants in human populations that may modulate its affinity to SARS-CoV-2 S-protein and thereby render individuals more resistant or susceptible to the virus. To investigate this, we shortlisted the *ACE2* non-synonymous variants fulfilling the triple requirement of 1) high allelic frequency (>10e-4 (gnomAD)), 2) involved critical residues in the ACE2-claw S-protein RBD-binding interface (29, 30) and 3) being potentially clinically relevant especially in Mediterranean populations (27, 28) (Figure 10A). Therefore we 5 shortlisted three common c.2158A>G p.(N720D), c.77A>G p.(K26R), and c.631G>A p.(G211R) SNPs and two rare variants, namely, c.1051C>G p.(L351V) and c.1166C>A p.(P389H) (Figure 10A). Intriguingly, these variants are overrepresented in the Mediterranean and European populations (*P* <0.001) but are extremely rare in the Asian population (27, 28, 31). We analyzed the impact of the shortlisted *ACE2* non-synonymous variants on SARS-CoV-2 infection using the Pseudovirus infection model described above. We found that upon similar expression of the *ACE2* variants (Supplemental Figure 1A), SARS-CoV-2 pseudovirus containing Wuhan-1 Spike was able to infect A549 cells more efficiently in K26R (*P*=0.0008), P389H (*P*=0.0012) and N720D (*P*=0.0059) as compared to the wild-type (WT) *ACE2* (Figure 10B). Conversely, we found that the variant G112R exhibited a moderate protective effect on infection (*P*=0.027). No significant differences were observed in L351V variant (p=0.768) compared to WT (Figure 10B). We obtained similar results when pseudotyping with Alpha, Beta, Delta or Zeta VOC (Figure 10C). This increased infectivity strongly suggests that bearing K26R, P389H or N720D *ACE2* variants could constitute a predisposing genetic background for severe COVID-19.

**Figure 10.**
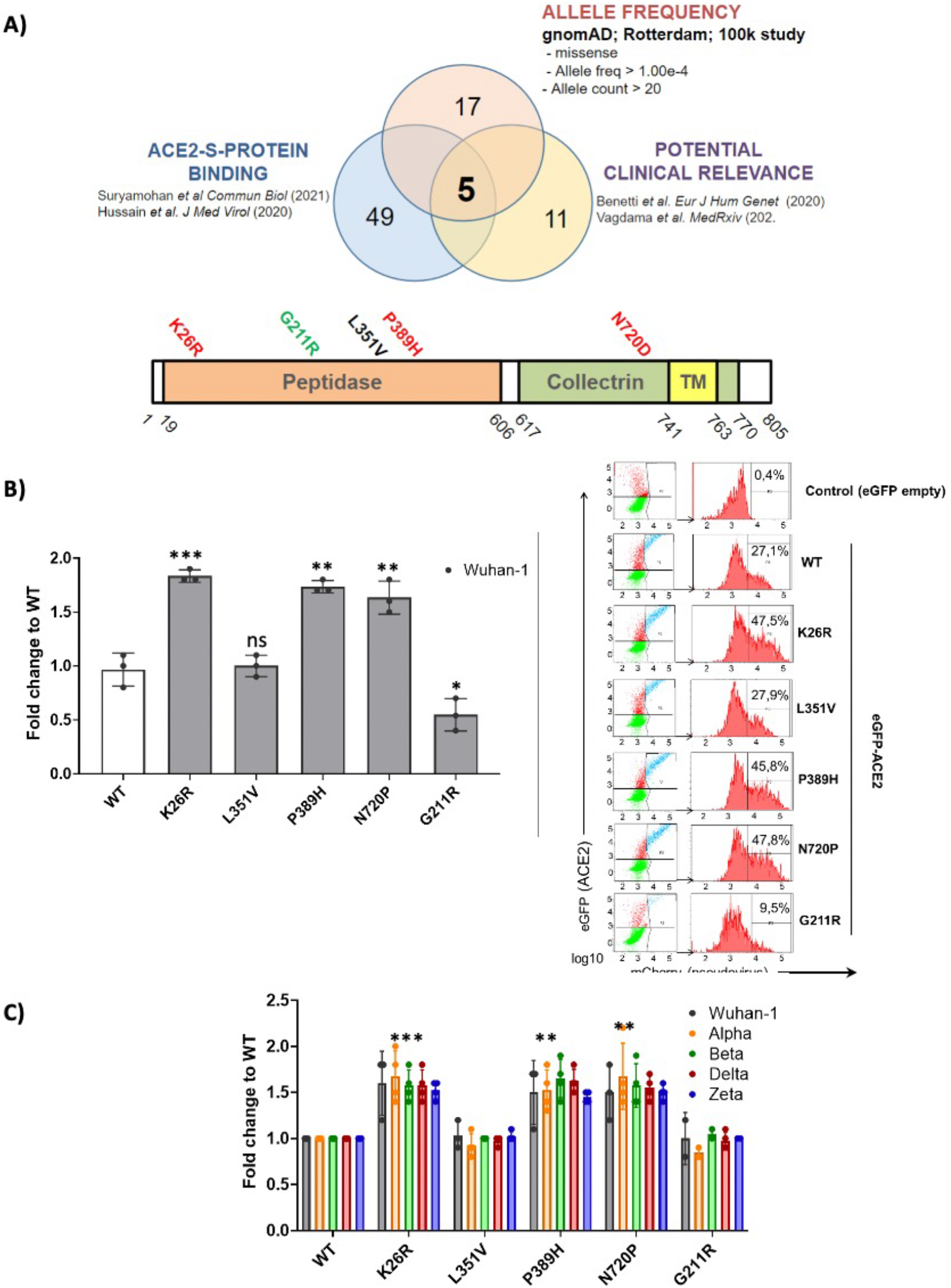
ACE2 polymorphisms K26R, P389 and N720D promote SARS-CoV-2 infection in all VOCs. A) Studying the effect of ACE2 SNPs. Non-synonymous ACE2 single nucleotide polymorphism were selected among those fulfilling the triple criteria of high allelic frequency (Allele freq > 1.00e-4; Allele count > 20); involved in ACE2-claw S-protein RBD-binding interface and previously associated to clinical outcome. B, C) Following the strategy described in Fig 9A, A549 cells expressing either GFP-ACE2 either WT or polymorphisms were assayed for cell entry by SARS-CoV-2 pseudovirus expressing either empty vector (Δ Spike) or Spike protein corresponding to origin variant (Wuhan-1) or variants-of-concern Alpha, Beta, Delta or Zeta. A representative flow cytometry experiment is shown. Bars demonstrate mean and Standard Error of Mean while each data point represents a unique experiment; ***P < 0.001, **P < 0.01, *P < 0.1, ns P >0.1 to WT by t test.

### 2.6. *ACE2* polymorphisms K26R, P389H and N720D are risk factor for severe COVID-19

Given the potential impact of shortlisted *ACE2* variants in SARS-CoV-2 infection and its overrepresentation in Mediterranean populations (27, 28), we analyzed the presence of these variants in severity-stratified groups from our cohort in order to get an insight on the role of *ACE2* variants on interindividual severity and susceptibility to COVID-19. We found a differential distribution among groups (Figure 11 and Table 2). Notably, and in line with our in vitro observations, we found that the “infection-promoting” variants K26R, P389H and N720D were enriched in COVID-positive (non-ICU and ICU groups) as compared to healthy controls (Fisher’s exact test *P* = 0.001; OR= 5.097; RR=3.439 at 95% CI) (Figure 11B and table 3), suggesting that they constitute a risk factor for COVID-19 patients. Conversely, “infection-protective” G211R variant had a modest protective effect (Fisher’s exact test *P* <0.001; OR=0.01327; RR= 0.2295 at 95% CI) (Table 3). While these associations need to be confirmed in larger sample sizes, they strongly suggest that these variants could play a central role in COVID-19 susceptibility.

**Figure 11.**
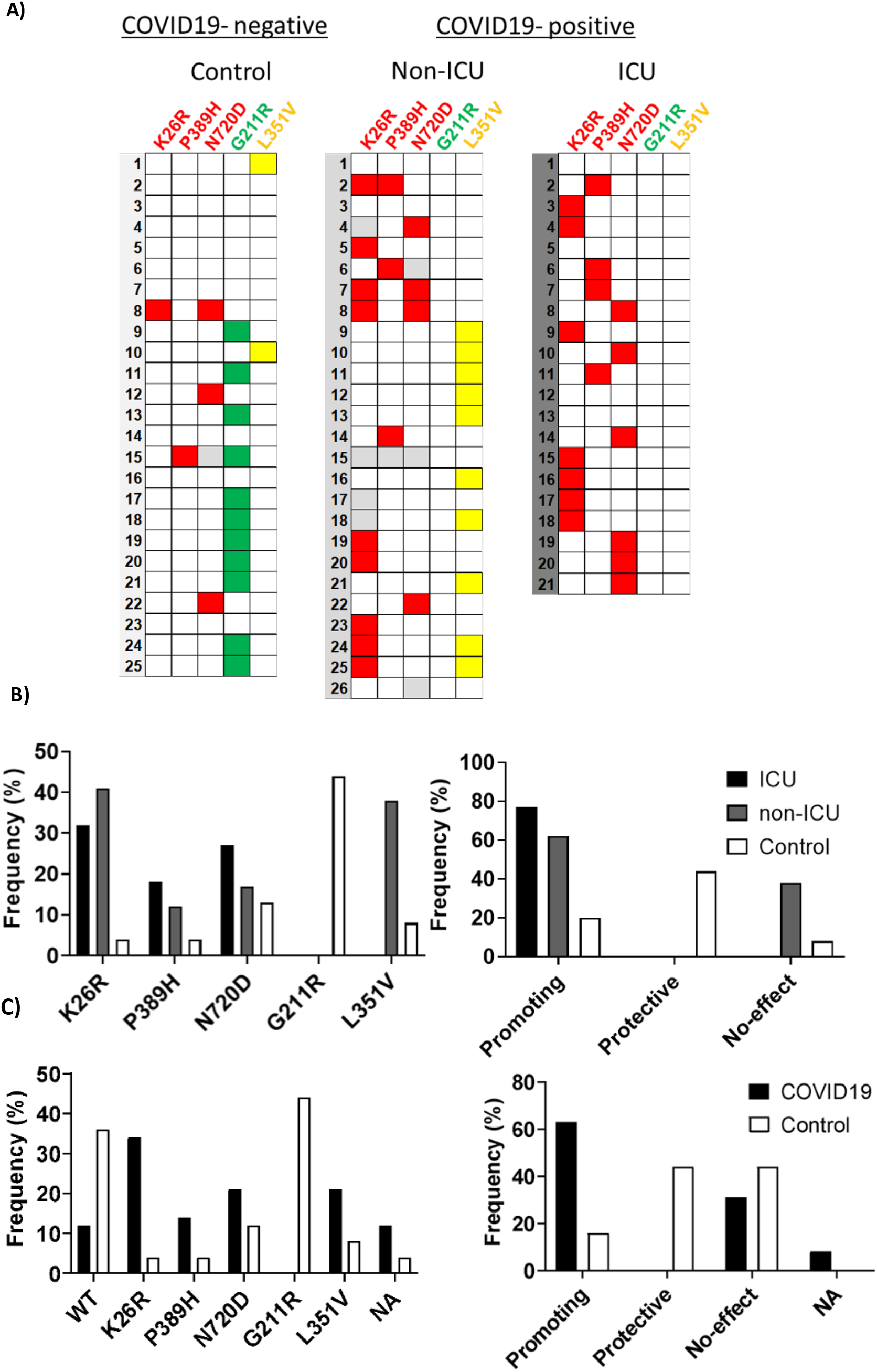
Infection-promoting ACE2 SNPs are a risk factor for COVID19 susceptibility. A) Heatmap showing the distribution of ACE2 variants in the hospitalization severity groups. Coloured squares indicate de presence of the ACE2 variants. Red: Promoting; Green: Protective; Yellow: No-effect. B) Frequencies of ACE2 SNPs among hospitalization severity groups. Bars represent frequencies of the SNP in each group C) Frequencies of ACE2 SNPS among susceptibility groups.

**Table 2.** *ACE2* SNP distribution of the sample population

|  | Total (n=72) | ICU<br>(n=21) | Non-ICU<br>(n=26) | Control group<br>(n=25) | Chi-square<br>*p*-value |
| --- | --- | --- | --- | --- | --- |
| **K26R** | 17/68 | 7/21 | 9/22 | 1/25 | 0.0081 |
| **P389H** | 8/71 | 4/21 | 3/25 | 1/25 | 0.2718 |
| **N720D** | 13/68 | 6/21 | 4/23 | 3/24 | 0.3795 |
| **G211R** | 11/72 | 0/21 | 0/26 | 11/25 | < 0.001 |
| **L351V** | 12/72 | 0/21 | 10/26 | 2/25 | < 0.001 |
| **Promoting** | 34/72 | 17/21 | 13/26 | 4/25 | 0.003 |
| **Protective** | 11/72 | 0/21 | 0/26 | 11/25 | < 0.001 |

**Table 3.** Infection-promoting *ACE2* polymorphisms are risk factor for COVID19 susceptibility in a 72 patient cohort.

|  | Healthy<br>(n=25) | COVID-19<br>patients<br>(n=47) | Relative risk<br>[95% CI] | Odds ratio<br>[95% CI] | Fisher's exact<br>test<br>*p*-value |
| --- | --- | --- | --- | --- | --- |
| **K26R** | 1/25 | 16/43 | 8.00<br>[1.168-54.79] | 14.22<br>[1.752-115.5] | 0.0028 |
| **P389H** | 1/25 | 7/46 | 3.048<br>[0.4741-19.59] | 4.308<br>[0.4985-37.23] | 0.2455 |
| **N720D** | 3/24 | 10/47 | 1.655<br>[0.5801-4.719] | 2.059<br>[0.5801-4.719] | 0.3553 |
| **G211R** | 11/25 | 0/47 | 0.2295 | 0.01327 | <0.0001 |
|  |  |  | [0.1449-0.3635] | [0.0007-0.2394] |  |
| **L351V** | 2/25 | 10/47 | 2.3 | 3.108 | 0.1958 |
|  |  |  | [0.6234-8.486] | [0.6242-15.48] |  |
| **Promoting** | 5/25 | 33/47 | 3.439 | 5.097 | 0.0014 |
|  |  |  | [1.346-8.790] | [1.719-15.11] |  |
| **Protective** | 11/25 | 0/47 | 0.2295 | 0.01327 | <0.0001 |
|  |  |  | [0.1449-0.3635] | [0.0007-0.2394] |  |

### 2.7. Influence of *ACE2* polymorphisms on inflammation markers

We then analyzed whether the different *ACE2* polymorphisms may influence the levels of inflammation markers. First, we tested if the presence or absence of the different variants could affect the levels of GDF15 or ACE2. Interestingly, we did not find differences in the circulating protein levels of GDF15 and ACE2 when taken the 72 samples as a whole (Supplemental Figure 3). However, the P389H, G211R, and L351V variants were associated with a decrease in the levels of circulating *ACE2* mRNA, while the individuals bearing the N720D variant showed increased levels of circulating *ACE2* mRNA (Supplemental Figure 4). Second, we repeated the analysis comparing ICU vs non-ICU group, and we found that the levels of GDF15 and ACE2 were similar in the ICU patients carrying the different mutations compared to the non-ICU group (Figure 12). Then, we decided to repeat the same analysis, but classifying our sample population in 4 genotypes based on the presence or absence of the different variants. Genotype 0 corresponding to subjects that did not carry any variant; Genotype 1, subjects that carry at least one promoting variant; Genotype 2, subjects that carry at least one protective variant; Genotype 3, subjects that carry at least one promoting and one protective variant (Supplemental Table 5; Figure 13). We only found differences in the levels of the *ACE2* mRNA (*P*<0.05).

**Figure 12.**
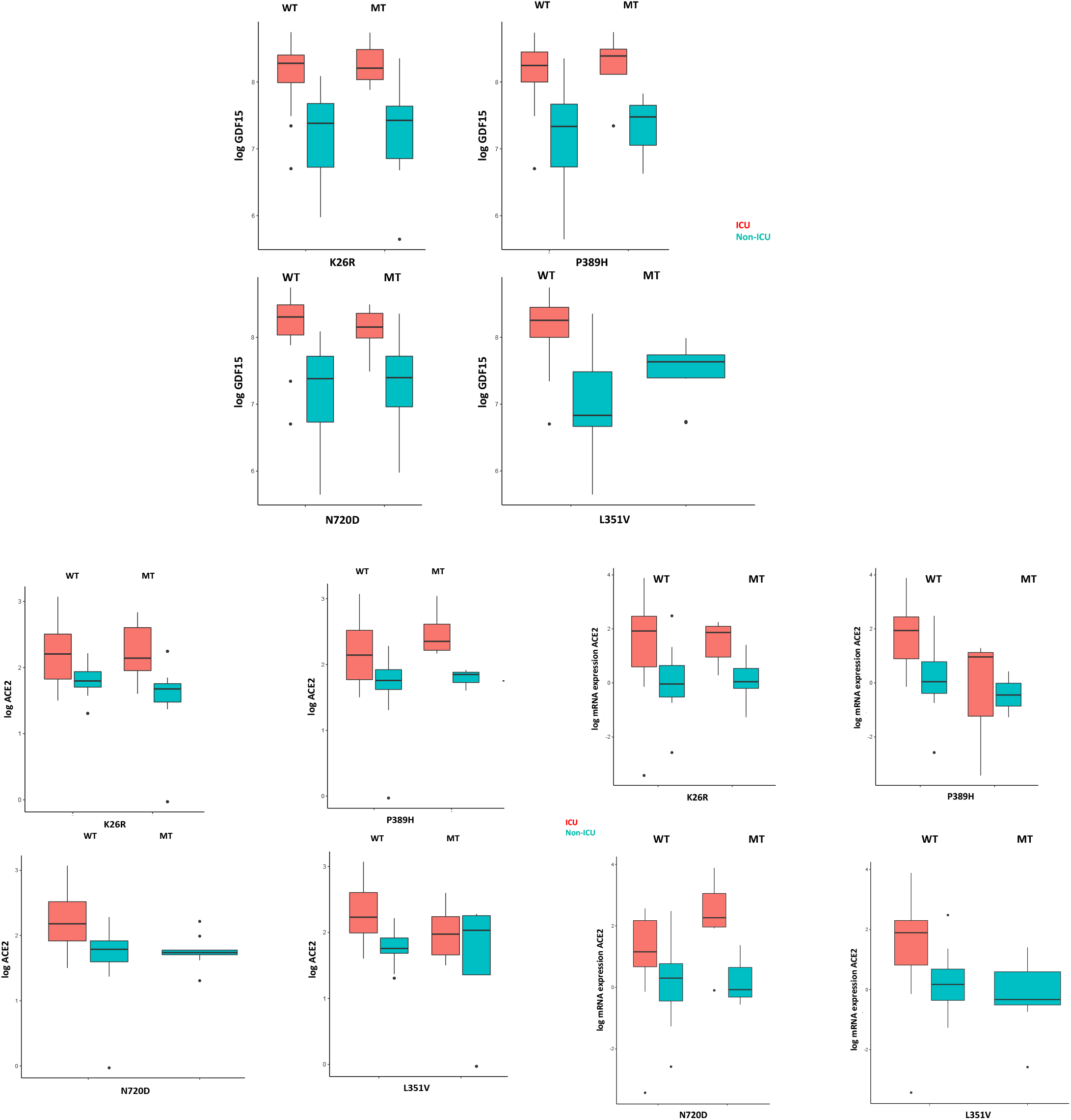
GDF15 and ACE2 levels among ICU and non-ICU COVID-19 patients and ACE2 genotypes.

**Figure 13.**
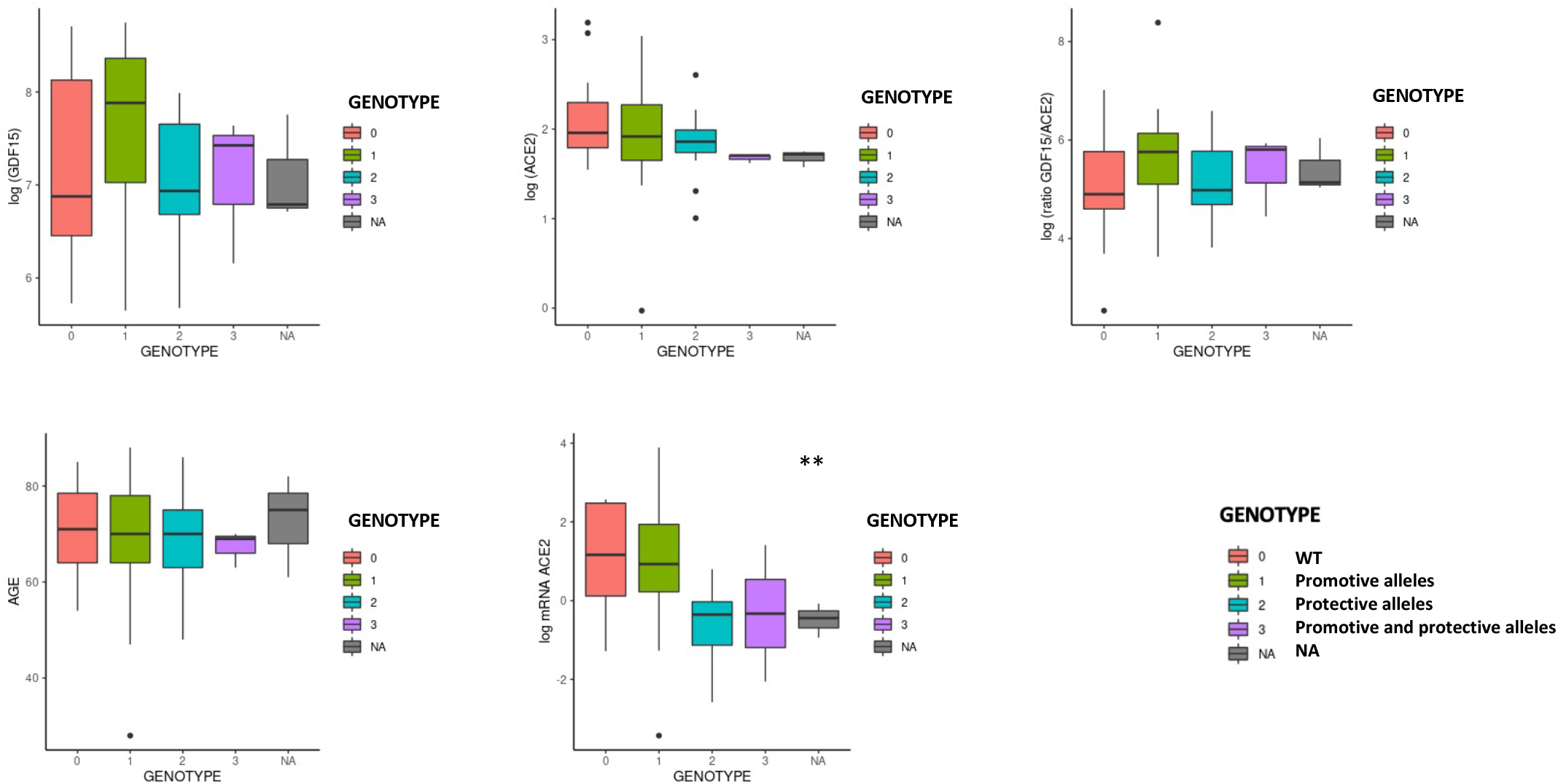
GDF15 and ACE2 levels by protective or promotive genotypes. Genotype 0 corresponding to subjects that did not carry any variant; Genotype 1, subjects that carry at least one promoting variant; Genotype 2, subjects that carry at least one protective variant; Genotype 3, subjects that carry at least one promoting and one protective variant (Supplemental Table 5; Figure 13). We only found differences in the levels of the ACE2 mRNA (*P*<0.05).

Finally, in linear regression models, adjusting for age, sex, and group (ICU, non-ICU, and Control group) we did not find any association between the *ACE2* genotypes with GDF15 and ACE2 levels (Supplemental Table 6) suggesting a separate role of these markers in COVID-19 susceptibility and severity.

## 3. Discussion

Despite the fact that the majority of the population is already vaccinated against SARS-CoV-2 and the rate of mortality has been reduced, vaccination is not enough to defeat the virus. COVID-19 infection rate remains high, and treatment of severe cases has become a great challenge. Therefore, early recognition of the sensitivity and severity of COVID-19 is essential for the development of new treatments. In the present study, we measured the levels of circulating GDF15 and ACE2 in plasma of patients with COVID-19 who were admitted to the ICU and died and admitted to the ICU who recovered, as well as in a healthy matched control group. We also characterized the in vitro effects and frequency of 5 *ACE2* SNPs overrepresented in Southern European populations and associated with COVID-19 disease (27, 28).

We found that both circulating GDF15 and ACE2 levels were higher in patients admitted to the ICU who died, compared to those who recovered from the disease and the control group. Our results are consistent with recent studies that have found that increased circulating levels of GDF15 and ACE2 are associated with a poor prognosis in patients with COVID-19 (11–13, 22, 23). On the other hand, we also found significant changes in *ACE2* gene expression levels in COVID-19 patients compared to healthy controls, as well as among COVID-19 patients. Previous studies have shown that ACE2 is highly expressed in infected tissues with SARS-CoV-2 compared to normal tissues (18, 32).

It is well established that both the occurrence and severity of COVID-19 increase with age, and the presence of comorbidities such as hypertension, diabetes, obesity, and cardiovascular disease conditions also largely related to the production of GDF15 (14, 16). In the present study, and as we expected, we found an association between the GDF15 levels and age while we did not find differences in ACE2 levels with age. Recently, Tanaka et al. characterized the plasma proteomic signature in humans in different age ranges and found that GDF15 had the strongest positive association with age (33). In addition, GDF15 was found to be one of the proteins in the secretory senescence-associated phenotype (SASP) protein repertoire, indicating the possibility that GDF15 modulates and/or predicts cellular senescence (34, 35).

Regarding gender, there were no differences in circulating GDF15 nor ACE2 levels in our studied population, although slightly higher levels were observed in men, but not statistically significant. Other studies have shown that ACE2 levels were higher in men than in women (36). Regarding the association of ACE2 levels and age, the data available in the literature are limited. AlGhatrif et al. found that ACE2 levels show a curvilinear association with age in healthy people, with a positive association among participants younger than 55 years and a negative association in participants older than 55 (37). A recent study found that ACE2 levels increase with age in patients with acute respiratory failure who required mechanical ventilation (38).

Elevated levels of classic inflammatory markers, such as CRP, ferritin, and D-dimer, are associated with a worse prognosis in patients with COVID-19 (39). We found that both, GDF15 and ACE2 levels, were positively correlated with those markers. In COVID-19, possible mechanisms behind systemic clinical findings include dysregulated iron homeostasis, resulting in oxidative stress and an inflammatory response. Dysregulation of iron homeostasis and higher iron levels may support the progression of viral infections (40). Also, it seems that GDF15 has a fundamental role in the regulation of iron metabolism through the modulation of hepcidin so that the deregulation of plasma levels of GDF15 could directly affect iron homeostasis (41). Our data are consistent with previous studies (23, 42). Interestingly, both circulating GDF15 and ACE2 were positively correlated with glucose, urea, creatinine, and NT-proBNP. GDF15 has been recently recognized as a metabolic regulator, due to its role in regulating appetite and metabolism and obesity through its receptor (43).

GDF15 is closely related to inflammation. GDF15 is a known marker of mitochondrial dysfunction, and due to its function as hormone-cytokine may play an important role in the regulation of cellular oxygenation and the inflammatory response, all of which are key mechanisms in the pathophysiology of COVID-19. GDF15 is upregulated in situations of hypoxia and tissue damage (44). However, knowledge of its pathophysiological function at the molecular level is still limited and more studies are needed. It has recently been discovered that the α-type glial cell-derived neurotrophic factor family receptor (*GFRAL*), expressed exclusively in the brain, could be key to the molecular mechanism of GDF15 and its relationship with pathological states (45). A recent study has shown that GDF15 can promote human rhinovirus-associated lung inflammation in mice, leading to severe respiratory viral infections (46). In contrast, GDF15 has also been proposed as a central mediator of tissue tolerance induced by inflammation, by protecting against bacterial and viral infections, as well as sepsis, in mouse models (47).

ACE2 is a membrane-bound enzyme and therefore measurement of circulating levels of ACE2 is complex. ACE2 is cleaved from the plasma membrane through ADAM17-mediated regulated removal (48). Therefore, elevated plasma ACE2 levels could also result from increased lysis of ACE2-expressing cells as a consequence of more severe COVID-19 infection (49). Contrary to what we expected, we found no association between the levels of GDF15 and the levels of ACE2 in plasma. However, a previous study GDF15 levels were positively associated with ACE2 levels in patients with atrial fibrillation (50).

Based on the all the above, we believe that both GDF15 and ACE2 could provide valuable information on the pathophysiology of COVID-19. More studies are needed to elucidate the role that these biomarkers play in the progression of SARS-CoV-2 infection.

Host genetics is known to play an important role in susceptibility to other viral infectious diseases, including SARS-CoV, HIV and influenza (51). The most well-known and evaluated mechanism of SARS-CoV-2 infection is the binding and uptake of viral particles through *ACE2* receptor, a type 1 integral membrane glycoprotein (52). Based on the premise that a predisposing genetic background may contribute to the wide clinical variability of COVID-19, we set out to investigate whether variations in ACE2 might modulate susceptibility to SARS-CoV-2 and influence severity. We queried multiple genomic databases comprising the main sources of aggregated data for ACE2 protein altering variations in populations groups across the world to considered non-synonymous allele variants with high allelic frequency (>1.00.e-4) and allele count >20. Then we selected variants within the human ACE2-claw S-protein RBD-binding interface as described in previous studies (29, 30) that could potentially increase or decrease the binding affinity of ACE2 to the S-protein and thereby alter the ability of the virus to infect the host cell. Then, we considered the ACE2 variants previously described potentially predisposing genetic background to the observed individual clinical variability (27, 28). Finally, five *ACE2* variants fulfilled this triple requirement. Notably, these variants have not been reported in the Asian population and are rare in non-South European populations (27, 28). Given their rarity in other populations, we cannot exclude that these variants can partially account for the clinical outcome observed in the Spanish population (27). Using a pseudovirus infection approach similar to the previously reported (53, 54) we recapitulated airway infection in human alveolar lung A549 cells. This cell line does not express detectable levels of ACE2, thus being an excellent model to test the effects of ectopic ACE2 containing particular SNPs. With this approach, we found that variants K26R, P389H and N720D promoted a significant increase in infection while G211R diminished the infection being potentially protective. Variant N720D lies in a residue located close to the cleavage sequence of TMPRSS2, likely affecting the cleavage dependent virion intake (29). Variant K26R was previously reported to increase affinity for S-protein by biochemical assays (29) while G211R was predicted in silico to affect S-protein and ACE2 interaction (27). In our model, variant L351V showed no significant difference in infection in contrast to the previous results by crystal structure (27).

We also studied the presence of the 5 specific SNPs in *ACE2* coding regions using human plasma circulating mRNA. Transcription of *ACE2* produces different mRNA transcript variants responsible for translation into different protein isoforms (55, 56) that could be regulated by COVID-19 infection in a cytokine-specific manner (57). Therefore, we reasoned that patient genotyping from mRNA could better reflect the *ACE2* variants expressed in those patients. In fact, previous studies have shown that SNPs could be detected with high precision in transcriptome sequencing approaches as compared to DNA-seq procedures (58, 59). This has led to the emergence of transcriptome or RNA sequencing as a potential alternative approach to variant detection within protein coding regions, since the transcriptome of a given tissue represents a quasi-complete set of transcribed genes (mRNAs) and other noncoding RNAs which also bypasses the need for exome enrichment (60, 61). Using this approach, we identified the presence of the 5 shortlisted SNPs involved in vitro in a differential susceptibility in the 72-patient cohort. Notably, infection-promoting variants K26R, P389H and N720D were a risk factor for Severe COVID-19 Hospitalized groups as compared to wild-type ACE2. Conversely, infection inhibiting variant G211R showed a protective effect as compared to wild-type ACE2. Interestingly G211R exhibited a gender bias, being significantly enriched in women. This is in agreement with the gender effect in COVID-19 susceptibility showed by others (25, 28).

Our data suggest that *ACE2* SNPs could stratify patients according to infection susceptibility in complete agreement with our pseudoviral infection model. Therefore, it is plausible to think that the effect of allelic variability on ACE2 conformation would at least partially account for the interindividual clinical differences and likely modulate clinical susceptibility. This finding reinforces the hypothesis that at least some of the identified variants or the cumulative effect of few of them could confer a different susceptibility to virus cell entry and consequently to disease onset.

Our study has limitations. First the study population is small, including patients from Hospital Son Llatzer and Hospital Son Espases, two of the main hospitals in the Balearic Health System. There might be a selection bias when identifying factors that differ between patients with COVID-19, although COVID-19 patients were matched for age and sex as well with the control group. Second, it is a retrospective study conducted in an emergency situation, and in which not all the clinical characteristics of the patients were recorded. Third, some patients had elevated biomarker data in some laboratory measurements, as well as in GDF15 and ACE2 levels, and we did not exclude them from the analysis due to small sample size and because those numbers were physiological and not due to technical errors. Finally, the study is cross-sectional and no causal inferences can be made. Despite the limitations, our study provides convincing evidence that in patients with COVID-19, the levels of GDF15 and ACE2 could be associated with increased inflammation and disease severity while *ACE2* missense SNPs might be linked to infection susceptibility.

In conclusion, critically ill patients with COVID-19 present higher levels of GDF15 and ACE2, as well as acute inflammation. These two proteins might be of importance because of its association with disease severity in patients infected with SARS-CoV-2. Our results suggest that certain genetic variations in *ACE2* might modulate susceptibility to SARS-CoV-2 infection and influence severity. More studies are needed to elucidate the role of GDF15 and ACE2 in COVID-19.

## 4. Methods

### 4.1 Patient population

This study included blood samples from 72 subjects: 47 COVID-19 samples were obtained from the Biobank Unit located in the Hospital Universitario Son Espases (HUSE), in Palma de Mallorca and 25 healthy patients (control group), matched by age and sex, obtained from the Blood and Tissue Bank of the Balearic Islands. COVID-19 patients were classified based on disease severity into two groups: those requiring intensive care unit (ICU) admission for more than 24 h and died (n=21), and those requiring only low care intensity hospitalization and recovered (n=26) (non-ICU). The SARS-CoV-2 infection was confirmed by a positive result of a real time RT-PCR from nasal or pharyngeal swabs. Blood and buffy coat samples were obtained from the patients and stored at −80 °C at the Biobank Unit. Clinical data of the individuals included in this study include age, sex, smoking habit, comorbidities, and inflammatory and coagulation markers (PCR, ferritin, and D dimer). Not all the subjects had complete clinical data available.

### 4.2. Determination of ACE2 and GDF15 circulating levels

Circulating levels of ACE2 from plasma samples were quantified using the Human ACE2 SimpleStep ELISA® Kit (#ab235649, Abcam) following the manufacturer’s instructions. Briefly, plasma samples were diluted 1:2 with the Sample Diluent NS and plated with the Antibody cocktail. Plates were incubated overnight at 4 °C with gentle rocking. The following day, plates were washed, the TMB Development Solution was incubated for 10 min and the Stop Solution was added. An endpoint reading was performed at 450 nm in the Biotek Synergy H1 microplate reader. All samples were assayed in duplicate.

GDF15 circulating levels were measured using the Human GDF15 Quantikine® ELISA Kit (#DGD150, R&D Systems). Plasma samples were diluted 1:4 with the Calibrator Diluent and plated and incubated overnight at 4 °C with gentle rocking. The next day, after washing the plates, Human GDF15 Conjugate was incubated by shaking for 1 h at room temperature. Then, the Substrate Solution was incubated for 30 min at room temperature in the dark and the Stop Solution was added. An endpoint reading was performed at 450 nm with a wavelength correction set at 540 nm in the Biotek Synergy H1 microplate reader. All samples were assayed in duplicate.

### 4.3. RNA and DNA isolation

RNA and DNA from plasma samples and buffy coats were extracted using TRI Reagent® BD (#T3809, Sigma-Aldrich) following the manufacturer’s protocol. RNA and DNA quantity and quality were evaluated using a Nanodrop 1000 spectrophotometer (Thermo Scientific) set at 260 and 280 nm. Samples were stored at −80 °C until further use.

### 4.4. RT-qPCR

For each sample, 200 ng of the total RNA isolated from plasma, or 1 µg of the total RNA extracted from the buffy coats were reversed transcribed to cDNA at 37 °C for 50 min. The reaction mixture contained 250 mM Tris-HCl (pH 8.3), 375 mM KCl, 15 mM MgCl2, 2.5 µM random hexamers, 500 µM of each dNTP, 20 U RNase inhibitor, 10 mM DTT, and 200 U M-Mlv reverse transcriptase. Each cDNA was diluted 1/10 with free RNAse water and frozen at −20 °C until further use. PCR reactions were performed on a LightCycler® 480 System. Genes, primers, and annealing temperatures are specified in Supplemental Table 1. Reaction mix contained 7.5 µL SYBR TB Green® Premix Ex Taq™ (RR420A, Takara) with 0.5 µM forward and reverse primers and 2.5 µL of each cDNA sample. The amplification program consisted of a denaturation step at 95 °C for 5 min, followed by 45 cycles with a denaturation step (10 s, 95 °C), an annealing step (10 s, temperature depending on primers), and an elongation step (12 s, 72 °C). A negative control was run in each assay. The relative expression levels of each gene were calculated using the ΔΔCt method and normalized to GAPDH expression.

### 4.5. Circulating mtDNA levels determination

Circulating levels of mtDNA were quantified using 5 ng of the isolated DNA from plasma samples. A mitochondrial gene (cytochrome C oxidase subunit III, *COX3*) and a nuclear gene (*GAPDH*) were amplified in a qPCR reaction. The reaction program for *COX3* amplification consisted of a denaturation step of 95 °C for 3 min, and 45 cycles of 95 °C for 10 s and 55 °C for 30 s. The levels of mtDNA were estimated using the ΔΔCt method and normalized to 18S expression. All primers are specified in Supplemental Table 1.

### 4.6. Analysis of mtDNA oxidation

Oxidation of mtDNA was performed as described previously in (62) using the DNA extracted from buffy coats. Briefly, 5 ng of total DNA were used to amplify two different regions of mtDNA which are sensitive to oxidative damage. Oxidation of the G nucleotide diminishes the efficiency of the PCR reaction. Oxidation levels were estimated with the Ct ratio of these two regions and normalized by mtDNA levels determined as reported before. All primers are specified in Supplemental Table 2.

### 4.7. Telomere length

Telomere length was analyzed as described by Joglekar et al (63) using the DNA isolated from buffy coats. Briefly, two PCR reaction were performed using 25 ng of total DNA, amplifying a telomere region and a single-copy gene (human beta globin, *HBG*). All primers are specified in Supplemental Table 1.

### 4.8 Generation of SARS-CoV-2 pseudovirus

*pcDNA3.1* plasmids encoding for Spike SARS-CoV-2 variants Alpha (B.1.1.7), Beta (B.1.351), Delta (B.1.617.2) and Epsilon (B.1.427/B.1.429) variants-of-concern (VOCs) were obtained from Dr. Thomas Peacock (Barclay lab, Imperial College London, UK). *psPAX2* was a gift from Dr Didier Trono (Addgene plasmid # 12260); *pLV-mCherry* was a gift from Dr Pantelis Tsoulfas (Addgene plasmid # 36084); *pcDNA3* mCherry was a gift from Dr Scott Gradia (Addgene plasmid # 30125); *pcDNA3.1*(+)eGFP was a gift from Dr Jeremy Wilusz (Addgene plasmid # 129020); *pcDNA3.1-*SARS2-Spike was a gift from Dr Fang Li (Addgene plasmid # 145032); *pcDNA3.1-ACE2*-GFP was a gift from Dr Utpal Pajvani (Addgene plasmid # 154962). *pcDNA3.1-*GFP-ACE2 expressing *ACE2* SNPs were synthesized by GenScript. To generate SARS-Cov-2 pseudovirus, Human embryonic kidney 293 (HEK 293, ATCC® CRL-1573™) cells were seeded in a 10 cm2-plate at 4.5×106 cells/plate, in DMEM (ThermoFisher Scientific, #11995073) supplemented with 10% FBS and 1x Normocin (Invivogen, #ant-nr-1) and grown overnight at 37 °C, 85% rel humidiy, 5% CO2. Next day, HEK 293 cells were then cotransfected with 15 μg of psPAX2 : pcDNA3.1SpikeVOCs : pLV-mCherry at ratio 1:1:1 (i.e. 5:5:5 μg) using Lipofectamin™ 3000 transfection reagent (Thermo-Fisher Scientific, #L3000015). Pseudoviruses harvested from the supernatant at 48 h 72h post-transfection were filtered (0.44 μm, Millipore Sigma, #SLHA033SS) concentrated with 50kDa-PES (ThermoFisher Scientific, #88540) at 20 min a 14,000g RT and stored at −80°C.

### 4.9. Pseudovirus entry assay

A549 cells (ATCC® CCL-185) seeded in 6-multiwell plates at 0.23 × 106 cells/well in DMEM (10% FBS and 1x Normocin), and grown overnight at 37 °C, 85% rel humidity 5% CO2. Next day, cells were transfected with either *pcDNA3.1*(+)eGFP (control) or *pcDNA3.1*-GFP-*ACE2* expressing the *ACE2* SNPs. To test the effect of *ACE2* SNPs on pseudovirus entry, at 24h posttrasfetion cells were infected with 0.25 mL concentrated pseudoviruses + 0.25 mL DMEM (10% FBS and 1x Normocin) overnight at 37 °C, 85% rel humidiy, 5% CO2. Next day, 1 ml of fresh DMEM was added (10% FBS and 1x Normocin). To measure the mCherry signal (a proxy for virus uptake), at 48h postinfection, medium was removed and cells were detached using TrypLE (ThermoFisher, #12605036) washed in DPBS (ThermoFisher, #14190250) and incubated for 5 min RT with 4ʹ,6-diamidino-2-phenylindole (DAPI) at a final concentration of 0.1 µg/mL. Viable (DAPI negative) double positive eGFP/mCherry cells were measured using a BD FACSAria Fusion cell Sorter (BD Bioscience). 10,000 total events were recorded for every sample. Samples with less than 5,000 total events were discarded. Flow cytometer data was analyzed using BD Bioscience Flow cytometry analysis software. Briefly, cell population was defined using FSH/SCC gating and the percentage of positive cells was calculated using a manually defined cutoff based on log10 fluorescence intensity histograms of the corresponding channel on a non-infected population.

### 4.10. Microscope imaging

In order to assess overall protein localization of *ACE2* variants, A549 cells grown at 40000 cells/well in 8-well cell culture chamber (Sarstedt, #94.6170.802) were transfected with 1,5 μL Lipofectamine 3000® (Thermo Scientific, #L3000015) following the manufacturer’s protocol. At 48h post transfection, cells were fixed with 4% paraformaldehyde in PBS for 10 minutes at room temperature (RT) and washed twice with PBS. Then, they were permeabilized with 0.1% Triton X-100-PBS for 15 min and incubated with 2% BSA in PBST (PBS+ 0.1% Tween 20) for 1 h at RT to block nonspecific binding. Chambers were incubated with either anti-ACE2 (1:100; MA5-32307; Thermo Scientific) or anti-ACE2 (1:50; MAB933; R&D Systems) in PBST containing 0,1% BSA ON at 4°C, washed twice in PBS and incubated with Alexa-Fluor647-labeled anti-rabbit (1:2000; ab150079; Abcam) or anti-mouse (1:2000; ab150115, Abcam) for 1h at RT. Images were acquired with Cell Observer-Zeiss.

### 4.11. Identification and selection of *ACE2* polymorphisms

We queried multiple genomic databases including gnomAD (Karczewski, et al.), RotterdamStudy (Ikram et al), UK’s 100k Genomes Project (100KGP) as the main sources of aggregated data for ACE2 protein altering variations in populations groups across the world. We only considered non-synonymous allele variants with high allelic frequency (>1.00.e-4) and allele count >20. Then we selected variants within the human ACE2-claw S-protein RBD-binding interface as described in previous studies that could potentially increase or decrease the binding affinity of ACE2 to the S-protein and thereby alter the ability of the virus to infect the host cell. Finally, we aimed to investigate which *ACE2* variants were associated with clinical outcome to shortlist the candidates to study in detail. Two different studies identified 5 *ACE2* variants as potentially predisposing genetic background to the observed interindividual clinical variability (27, 28). We finally shortlisted the 5 variants fulfilling this triple requirement of being abundant, involving critical residues in RBD interface and being potentially clinically relevant.

### 4.12. *ACE2* polymorphism genotyping

cDNA obtained from circulating mRNA was analyzed to detect selected *ACE2* SNPs. Three different PCR reactions were designed to cover the regions comprising the studied *ACE2* SNPs. Primers were designed to obtain amplicons of 0.2-0.8 kb and specificity was confirmed by PrimerBLAST (64). PCRs were performed with GoTaq® G2 Flexi DNA Polymerase (Promega; M7805). PCR conditions were optimized by the Promega PCR design software (https://www.promega.es/resources/tools/biomath/tm-calculator/). PCR1 amplified the region corresponding to residues Ser3 to Met249 (743 bp). Primers Forward 5’AAGCTCTTCCTGGCTCCTTC3’, Reverse 5’TCATCAACTTTGCCCTCACA3’. Cycling conditions were as follows: 95°C for 5 minutes, followed by 40 cycles (94°C for 30s, 67°C for 30s, 72°C for 1 min) with a final extension at 72°C during 10 min.

PCR2 amplified the region corresponding to residues Phe308 to Arg621 (944 bp). Primers Forward 5’ ATTCAAGGAGGCCGAGAAGT3’, Reverse 5’TCCTCACTTTGATGCTTTGG3’. Cycling conditions were as follows: 95°C for 5 minutes, followed by 40 cycles (94°C for 30s, 63°C for 30s, 72°C for 1 min) with a final extension at 72°C during 10 min.

PCR3 amplified the region corresponding to residues Val670 to Val752 (250 bp). Primers Forward 5’ TGTGCGAGTGGCTAATTTGA3’, Reverse 5’ CACTCCCATCACAACTCCAA3’. Cycling conditions were as follows: 95°C for 5 minutes, followed by 40 cycles (94°C for 30s, 65°C for 30s, 72°C for 30s) with a final extension at 72°C during 10 min.

Agarose gel electrophoresis was performed in order to confirm specificity. PCR product was purified by clean-up (FAGCK001, Favorgen) and further sequenced by Macrogen. Alignment of sequencing reads against reference *ACE2* (NM_021804) and SNP genotyping was attained by the SnapGene® Software (snapgene.com). Supplemental Figure 2

### 4.13. Statistical analysis

Descriptive characteristics were summarized as means and standard deviations (SDs) or as numbers and percentages (%). One-way analysis of variance (ANOVA), Chi-square tests (χ2), and Fisher’s exact test were used to assess differences across 3 groups of participants: ICU; non-ICU and controls, for continuous and categorical variables respectively. Non-normally distributed continuous data were compared using the Mann-Whitney-Wilcoxon test or the Kruskal-Wallis H-test. The relationships between variables were studied using Spearman or Pearson correlations Linear regression modeling was used to quantify the associations of *ACE2* genotypes on GDF15 and ACE2. Analyses were adjusted for: age, sex, and group. Correlation coefficients were performed to determine correlations between GDF15 and changes on biochemical parameters among COVID-19 infected patients. To test potential associations between the studied parameters, a Principal Component Analysis (PCA) was computed. Statistical analyses were performed using R Studio version 3.5.2 of the R programming language (R Project for Statistical Computing; R Foundation, Vienna, Austria). and Stata v17.0 program. *P*-values <0.05 were deemed as statistically significant.

### 4.14 Study approval

This study was conducted in agreement with the Good Clinical Practice principles and the Declaration of Helsinki for ethical research. Ethical approval for this project (IB 4165/20 PI, 6 April 2020) was obtained from the ethics committee of the Balearic Islands and waived the requirement for informed consent, due to the emergency situation.

## 5. Author contributions

Conceptualization and methodology, MGF and CB; MTM, CMPR, NTF, LIG, AMGP, CNE and CB performed experiments, analyzed the data and wrote—original draft preparation; LSC, ASP, JPL, CC and LM collected and contributed data. Writing—review and editing, MGF, CB, MTM, CMPR, ASP and L.M.; supervision, MGF and CB; Project administration and funding acquisition, MGF and CB. All authors have read and agreed to the published version of the manuscript. MTM and CMPR share co-first authorship; MGF and CB are the co-corresponding authors.

## 6. Acknowledgments

We would like to thank Dr Gabriel Bretones and Prof Carlos López-Otín (Universidad de Oviedo, Spain) for kindly providing psPAX2 pLV-mCherry vectors and protocol for pseudovirus production. We thank Dr Thomas Peacock and Prof Wendy Barclay (Imperial College London, UK) for kindly providing SARS-CoV-2 VOC vectors.

This research was supported by the Miguel Servet program (MS19/00201 and MS19/00100) (MGF and CB), Instituto de Salud Carlos III (ISCIII), Madrid; Fondo extraordinario COVID —ISCIII (COV20/00571) (CB); Fundació LaMaratóTV3 (167-C-2021 51); the Margalida Comas Program (PD/050/2020), Comunidad Autonoma de las Islas Baleares (MTM); the FOLIUM fellowship program (FOLIUM 19/01), Impost turisme sostenible/Govern de les Illes Balears (AMPG and CMPR); and the TECH fellowship program, Impost turisme sostenible/Govern de les Illes Balears (TECH19/03) (LYG).

## Supplemental material

**Supplemental figure 1.**
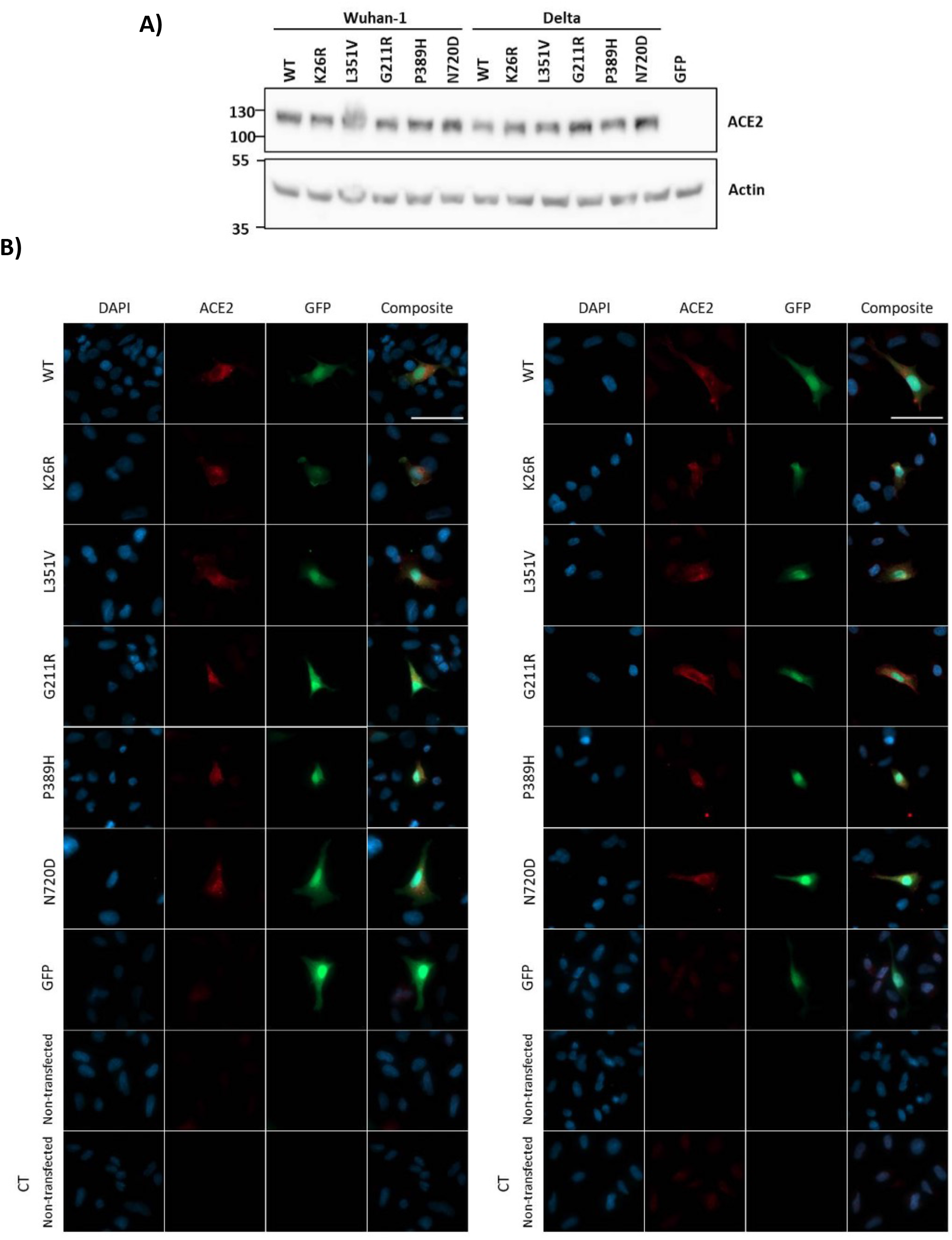
ACE2 polymorphisms exhibit similar fashion when expressed in A549 cells. A549 cells were transfected with either GFP-ACE2 WT, GFP-ACE2 polymorphisms or GFP alone. Then, ACE2 protein expression was analyzed by A) Western Blot with MA5-32307 antibody B) Immunocytochemistry (red) with either MA5-32307 antibody (left panel) or MAB933 antibody (right panel). Nuclei was stained with DAPI (blue). Transfected cells contain GFP (green). CT: secondary antibody control to detect unspecific binding. Images were acquired with Cell Observer-Zeiss. Scale bar: 50 μm

**Supplemental figure 2.**
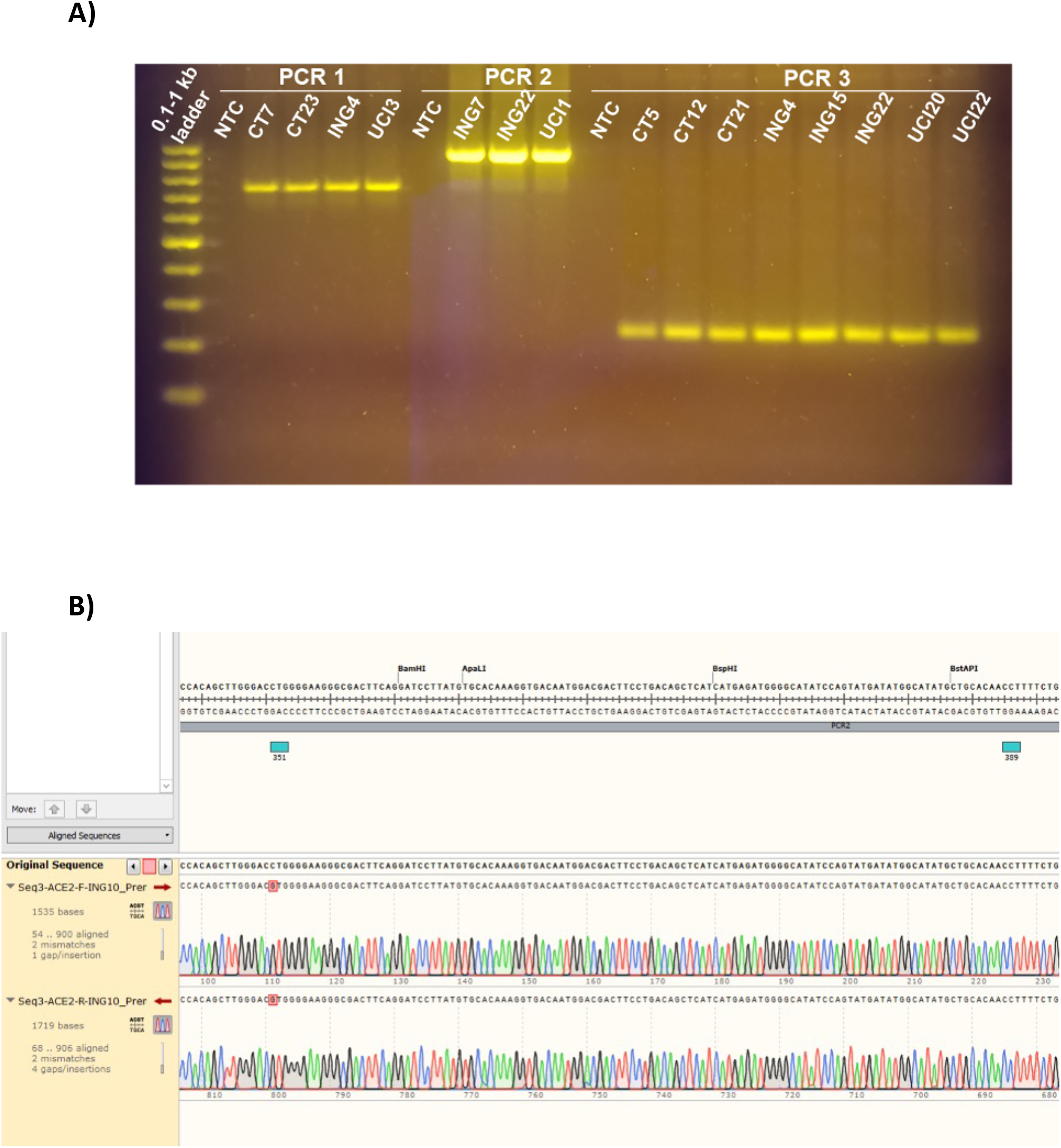
ACE2 missense SNPs genotyping. cDNA was obtained by RT-PCR from circulating mRNA of the patients. Then, PCR1, PCR2 and PCR3 were performed in order to amplify the regions comprising the studied SNPs. PCR products were sequenced and aligned against reference ACE2 (NM_021804). PCR1: residues from Ser3 to Met249 (743 bp), PCR2: residues Phe308 to Arg621 (944 bp), PCR3: residues from Val670 to Val752 (250 bp) A) Agarose gel electrophoresis with the PCR products of several patients was performed to confirm specificity. B) Representative image of the alignment of the sequenced (forward and reverse) PCR products against reference *ACE2* using SnapGene® Software. In particular, the image corresponds to the PCR2 of the non-ICU patient 10 that presents the L351V variant.

**Supplemental Figure 3.**
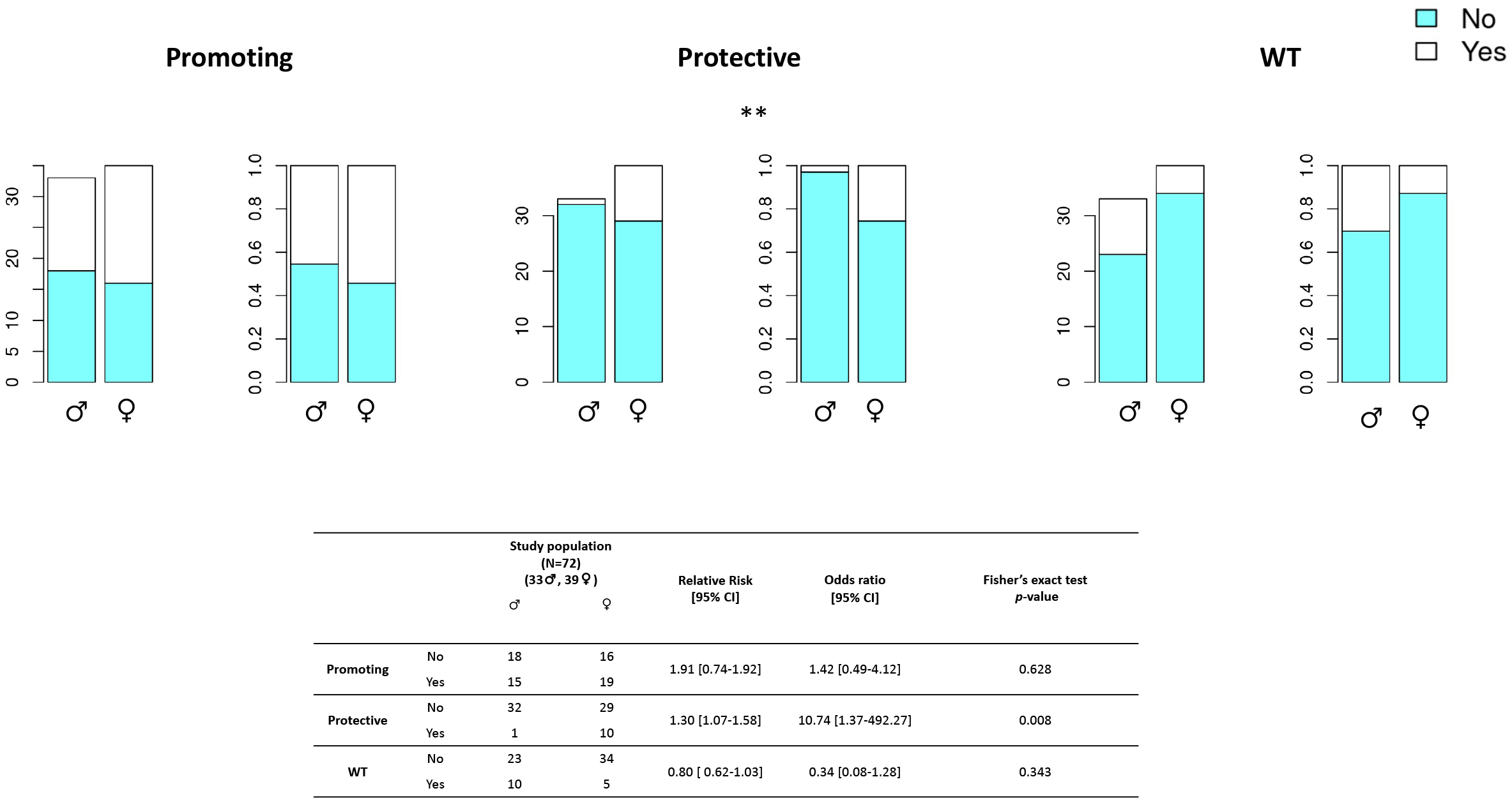
Frequency distribution of *ACE2* genotypes by sex. A) ‘promoting’ alleles; B) ‘Protective’ alleles: C) ‘WT’ alleles.

**Supplemental Figure 4.**
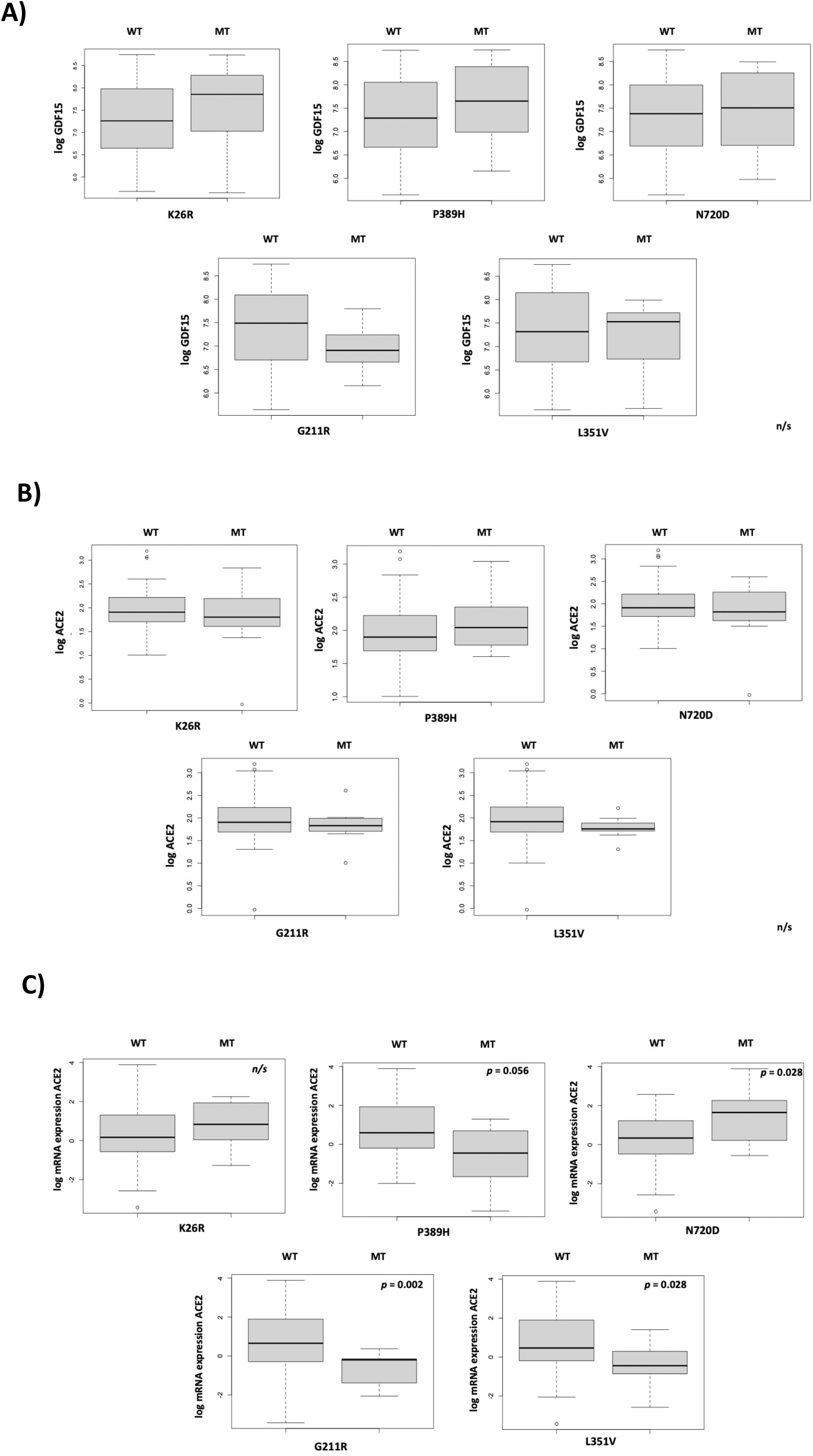
GDF15 and ACE2 levels across *ACE2* variants.

**Supplemental Table 1.**
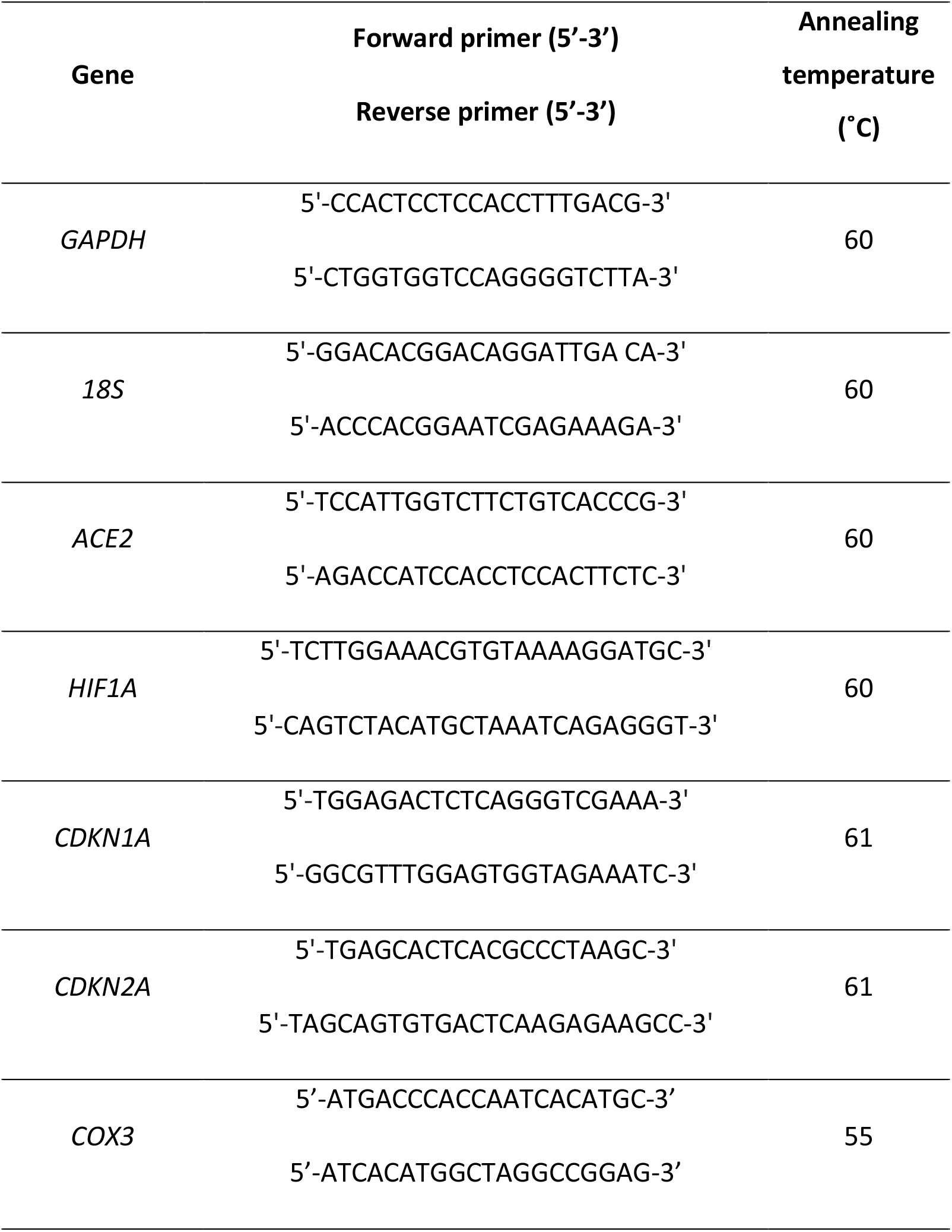
Primer sequences and annealing temperatures.

**Supplemental Table 2.**
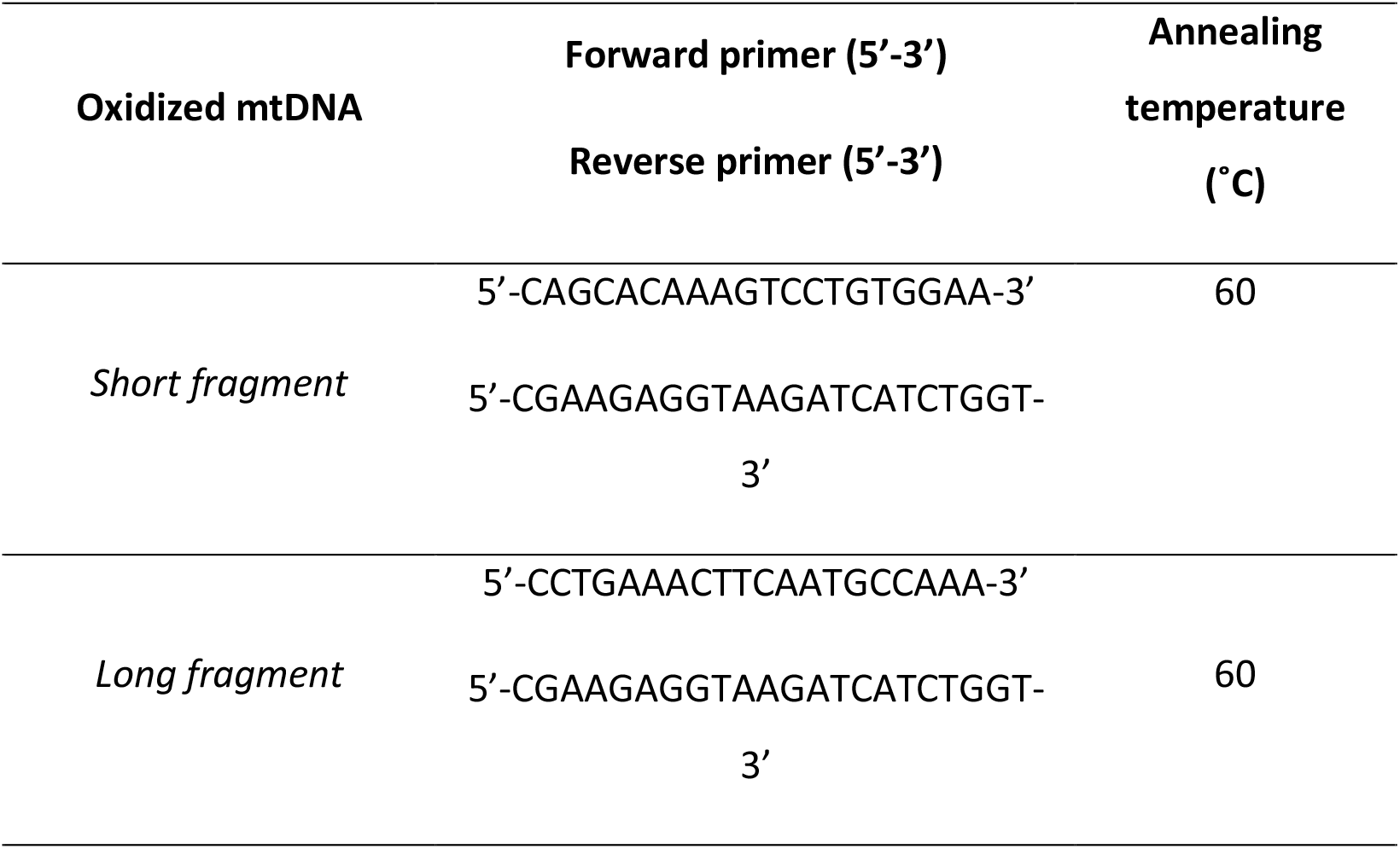
Primer sequences and annealing temperatures for assessment of mtDNA oxidation.

**Supplemental Table 3.** Clinical and Biochemical Characteristics of non-ICU group

| **Sociodemographic and history illness information** | No-ICU group, n=21 |  |
| --- | --- | --- |
|  | Mean | SD |
| Age (years) | 71.3 | 7.28 |
| Sex (women), n (%) | 9 | 42.9 |
| Smoking habit (Yes), n (%) | 13 | 61.9 |
| Hypertension (Yes), n (%) | 13 | 61.9 |
| T2DM (Yes), n (%) | 10 | 47.6 |
| Obesity (Yes), n (%) | 2 | 9.5 |
| COPD (Yes), n (%) | 0 | 0 |
| Dyslipidaemia (Yes), n (%) | 10 | 47.6 |
| **Biochemical parameters** |  |  |
| GDF15 | 1557 | 852 |
| ACE2 | 6.43 | 1.75 |
| Leukocytes, 10<sup>9</sup>/L | 6.99 | 2.01 |
| Neutrophil count, 10<sup>9</sup>/L | 4.26 | 1.73 |
| Lymphocyte count, 10<sup>9</sup>/L | 1.97 | 0.67 |
| Monocyte count, 10<sup>9</sup> /L | 0.57 | 0.19 |
| Eosinophils, , 10<sup>9</sup> /L | 0.16 | 0.09 |
| Haematids | 4.45 | 0.55 |
| Hemoglobin, g/dL | 13.7 | 1.45 |
| Hematocrit, % | 40.7 | 4.56 |
| Mean Corpuscular Volume (MCV), f | 91.8 | 4.60 |
| Mean Corpuscular Hemoglobin (MCH), pg | 31.1 | 2.15 |
| Red Blood Cell Distribution Width (RDW), % | 13.1 | 1.02 |
| Platelets, 10<sup>9</sup>/L | 206 | 48.7 |
| Mean Platelet Volume (MPV), f | 8.48 | 1.23 |
| Platelet Distribution width (PDW), % | 16.1 | 0.66 |
| Neutrophils - Lymphocytes ratio | 2.41 | 1.44 |
| Protombin ratio | 1.27 | 0.96 |
| D-Dimer, ng/mL | 172 | 177 |
| Glucose, mg/dL | 124 | 64.7 |
| Urea, mg/dL | 45.6 | 19.1 |
| Creatinine, mg/dL | 1.02 | 0.48 |
| Aspartate aminotransferase (AST), U/L | 19 | 4.31 |
| Alanine aminotransferase (ALT), U/L | 17.3 | 8.16 |
| Gamma-glutamyl transferase (GGT), U/L | 22.3 | 10.6 |
| Lactate dehydrogenase (LDH), U/L | 204 | 31.3 |
| Ferritin, ng/mL | 108 | 72.2 |
| C reactive protein (CRP), mg/L | 0.63 | 0.97 |
| Triglycerides, mg/dL | 132 | 49.8 |
| Nt PRO BNP, pg/mL | 385 | 868 |

Data shown is mean (SD), unless otherwise specified. The sample of the present analyses was n=21, corresponding to the non-ICU group. Abbreviations: SD: Standard deviation; ACE2: angiotensin 2-converting enzyme; GDF15: growth differentiation factor 15; T2DM: Type 2 diabetes mellitus; COPD: chronic obstructive pulmonary disease; Hb: haemoglobin; VCM: mean corpuscular volume; PDW VPM: mean platelet volume; LDH: lactate dehydrogenase; GGT: gamma glutamyl transpeptidase; ALT: alanine aminotransferase; AST: aspartate aminotransferase; CPR: c-reactive protein; NT\_PROBNP: brain natriuretic peptide.

**Supplemental Table 4.** Correlation coefficient between GDF15, ACE2 and changes on biochemical parameters among non-ICU COVID19 patients.

|  | GDF15 | ACE2 | Age | Ferritin | CRP | ACE2<sup>2-ΔΔCt</sup> | HIF1A<sup>2-ΔΔCt</sup> | ACE2<sup>2-ΔΔCt#</sup> | HIF1A<sup>2-ΔΔCt#</sup> | P16<sup>2-ΔΔCt#</sup> | P21<sup>2-ΔΔCt#</sup> | mtDNA/<br>nDNA<sup>#</sup> | mtDNAox<br>2-ΔΔCt# | TL<br>2-ΔΔCt# |
| --- | --- | --- | --- | --- | --- | --- | --- | --- | --- | --- | --- | --- | --- | --- |
| **GDF15** | 1.000 |  |  |  |  |  |  |  |  |  |  |  |  |  |
| **ACE2** | 0.304 | 1.000 |  |  |  |  |  |  |  |  |  |  |  |  |
| **Age** | **0.450\*** | -0.026 | 1.000 |  |  |  |  |  |  |  |  |  |  |  |
| **Ferritin** | **0.480\*** | 0.391 | -0.209 | 1.000 |  |  |  |  |  |  |  |  |  |  |
| **CRP** | 0.281 | 0.334 | -0.149 | **0.482\*** | 1.000 |  |  |  |  |  |  |  |  |  |
| **ACE2<sup>2-ΔΔCt</sup>** | 0.395 | 0.370 | -0.281 | **0.862\*\*\*** | 0.330 | 1.000 |  |  |  |  |  |  |  |  |
| **HIF1A<sup>2-ΔΔCt</sup>** | -0.163 | 0.021 | -0.330 | -0.128 | -0.162 | -0.177 | 1.000 |  |  |  |  |  |  |  |
| **ACE2<sup>2-ΔΔCt#</sup>** | 0.102 | 0.273 | -0.397 | 0.184 | -0.052 | 0.157 | **0.704\*\*** | 1.000 |  |  |  |  |  |  |
| **HIF1A<sup>2-ΔΔCt#</sup>** | -0.262 | 0.104 | -0.041 | -0.161 | -0.259 | **0.637\*\*** | -0.257 | -0.062 | 1.000 |  |  |  |  |  |
| **P16<sup>2-ΔΔCt#</sup>** | 0.118 | -0.657 | -0.473 | -0.532 | 0.086 | 0.602 | 0.578 | 0.810 | -0.041 | 1.000 |  |  |  |  |
| **P21<sup>2-ΔΔCt#</sup>** | -0.237 | -0.016 | -0.256 | -0.119 | -0.134 | -0.124 | **0.656\*** | **0.590\*** | 0.087 | **0.983\*\*** | 1.000 |  |  |  |
| **mtDNA/**<br>**nDNA** | -0.178 | 0.056 | -0.121 | -0.187 | **0.636\*\*** | -0.196 | 0.109 | -0.218 | -0.091 | 0.225 | 0.024 | 1.000 |  |  |
| **mtDNAox**<br>**2-ΔΔCt#** | **-0.495\*** | 0.040 | -0.349 | -0.128 | 0.196 | -0.138 | 0.287 | -0.050 | 0.184 | -0.238 | 0.326 | **0.576\*\*** | 1.000 |  |
| **TL<sup>2-ΔΔCt#</sup>** | -0.271 | 0.012 | -0.271 | -0.172 | **0.488\*** | -0.168 | 0.057 | 0.175 | -0.048 | 0.313 | 0.245 | **0.590\*\*** | 0.317 | 1.000 |

|  | **GDF-15** | **ACE2** | **ACE2<sup>2-</sup><sub>ΔΔCt</sub>** | **ACE2<sup>2-</sup><sub>ΔΔCt#</sub>** | **HIF1A<sup>2-</sup><sub>ΔΔCt</sub>** | **HIF1A<sup>2-</sup><sub>ΔΔCt#</sub>** | **Leukocytes** | **Neutrophil** | **Lymphocyte** | **Monocyte** | **Eosinophils** | **Basophils** | **Platelets** | **Neutro to**<br>**Lymp ratio** |
| --- | --- | --- | --- | --- | --- | --- | --- | --- | --- | --- | --- | --- | --- | --- |
| **GDF-15** | 1.000 |  |  |  |  |  |  |  |  |  |  |  |  |  |
| **ACE2** | 0.304 | 1.000 |  |  |  |  |  |  |  |  |  |  |  |  |
| **ACE2<sup>2-</sup><sub>ΔΔCt</sub>** | 0.395 | 0.370 | 1.000 |  |  |  |  |  |  |  |  |  |  |  |
| **ACE2<sup>2-</sup><sub>ΔΔCt#</sub>** | 0.102 | 0.273 | 0.157 | 1.000 |  |  |  |  |  |  |  |  |  |  |
| **HIF1A<sup>2-</sup><sub>ΔΔCt</sub>** | -0.163 | 0.021 | -0.177 | **0.704\*\*** | 1.000 |  |  |  |  |  |  |  |  |  |
| **HIF1A<sup>2-</sup><sub>ΔΔCt#</sub>** | -0.262 | 0.104 | **0.637\*** | -0.062 | -0.257 | 1.000 |  |  |  |  |  |  |  |  |
| **Leukocytes** | 0.163 | 0.396 | 0.363 | 0.189 | -0.112 | 0.067 | 1.000 |  |  |  |  |  |  |  |
| **Neutrophil** | 0.242 | 0.394 | **0.478\*** | 0.274 | -0.071 | 0.068 | **0.929\*\*\*** | 1.000 |  |  |  |  |  |  |
| **Lymphocyte** | -0.197 | 0.023 | -0.237 | -0.203 | -0.155 | 0.103 | 0.378 | 0.022 | 1.000 |  |  |  |  |  |
| **Monocyte** | 0.204 | 0.329 | 0.201 | -0.019 | -0.096 | -0.259 | **0.611\*\*** | **0.528\*** | 0.159 | 1.000 |  |  |  |  |
| **Eosinophils,** | 0.169 | 0.436 | -0.082 | **0.542\*** | 0.388 | 0.154 | 0.250 | 0.171 | 0.119 | 0.212 | 1.000 |  |  |  |
| **Basophils,** | -0.400 | -0.042 | -0.273 | -0.149 | -0.151 | -0.204 | 0.337 | 0.220 | **0.433\*** | -0.042 | -0.128 | 1.000 |  |  |
| **Platelets,** | **-0.557\*\*** | -0.159 | -0.261 | 0.026 | 0.124 | -0.426 | 0.289 | 0.079 | **0.576\*\*** | 0.276 | -0.091 | **0.479\*** | 1.000 |  |
| **Neutro to**<br>**Lymp ratio<sup>1</sup>** | 0.413 | 0.302 | **0.629\*\*** | 0.339 | -0.028 | -0.077 | **0.500\*** | **0.750\*\*\*** | **-0.530\*** | 0.341 | -0.004 | -0.078 | -0.329 | 1.000 |

| | GDF15 | ACE2 | ACE2<br>2-$\Delta\Delta$Ct | ACE2<br>2-$\Delta\Delta$Ct# | HIF1A<br>2-$\Delta\Delta$Ct | HIF1A<br>2-$\Delta\Delta$Ct# | mtDNAox<br>2-$\Delta\Delta$Ct# | mtDNA/<br>nDNA# | Glucose | Urea | Creatinine | TG | Nt_PRO<br>BNP | Prothrombi<br>n time |
| --- | --- | --- | --- | --- | --- | --- | --- | --- | --- | --- | --- | --- | --- | --- |
| **GDF15** | 1.000 |  |  |  |  |  |  |  |  |  |  |  |  |  |
| **ACE2** | 0.304 | 1.000 |  |  |  |  |  |  |  |  |  |  |  |  |
| **ACE2<sup>2-$\Delta\Delta$Ct</sup>** | 0.395 | 0.370 | 1.000 |  |  |  |  |  |  |  |  |  |  |  |
| **ACE2<sup>2-$\Delta\Delta$Ct#</sup>** | 0.102 | 0.273 | 0.157 | 1.000 |  |  |  |  |  |  |  |  |  |  |
| **HIF1A<sup>2-$\Delta\Delta$Ct</sup>** | -0.163 | 0.021 | -0.177 | **0.704\*\*** | 1.000 |  |  |  |  |  |  |  |  |  |
| **HIF1A<sup>2-$\Delta\Delta$Ct#</sup>** | -0.262 | 0.104 | **0.637\*** | -0.062 | -0.257 | 1.000 |  |  |  |  |  |  |  |  |
| **mtDNAox**<br>**2-$\Delta\Delta$Ct#** | -0.348 | -0.107 | -0.201 | -0.059 | -0.003 | -0.176 | 1.000 |  |  |  |  |  |  |  |
| **mtDNA/**<br>**nDNA#** | -0.178 | 0.056 | -0.196 | -0.218 | 0.109 | -0.091 | 0.149 | 1.000 |  |  |  |  |  |  |
| **Glucose** | **0.526\*** | 0.398 | **0.839\*\*\*** | 0.191 | -0.189 | **0.582\*** | -0.279 | -0.318 | 1.000 |  |  |  |  |  |
| **Urea** | **0.727\*\*** | 0.309 | **0.608\*\*** | 0.095 | -0.155 | -0.110 | -0.411 | -0.245 | **0.633\*\*** | 1.000 |  |  |  |  |
| **Creatinine** | **0.705\*\*** | 0.310 | **0.736\*\*** | 0.010 | -0.142 | 0.120 | -0.345 | 0.049 | **0.706\*\*** | **0.813\*\*\*** | 1.000 |  |  |  |
| **TG** | **0.486\*** | 0.143 | 0.071 | -0.196 | -0.235 | -0.144 | 0.351 | -0.224 | 0.148 | 0.268 | 0.270 | 1.000 |  |  |
| **Nt_PRO**<br>**BNP** | **0.494\*** | 0.351 | **0.889\*\*\*** | 0.176 | -0.126 | -0.248 | -0.217 | -0.224 | **0.944\*\*\*** | **0.665\*\*** | **0.784\*\*\*** | 0.161 | 1.000 |  |
| **Prothrombin**<br>**time** | 0.172 | 0.024 | -0.177 | -0.086 | 0.075 | -0.131 | -0.256 | **0.628\*\*** | -0.106 | -0.088 | 0.159 | -0.167 | -0.028 | 1.000 |

|  | **GDF15** | **ACE2** | **ACE2**<br>2<sup>-ΔΔCt</sup> | **ACE2**<br>2<sup>-ΔΔCt#</sup> | **HIF1A**<br>2<sup>-ΔΔCt</sup> | **HIF1A**<br>2<sup>-ΔΔCt#</sup> | **AST** | **ALT** | **GGT** | **LDH** | **D-Dimer** | **HCT** | **Hb** | **RBC** | **Igg** |
| --- | --- | --- | --- | --- | --- | --- | --- | --- | --- | --- | --- | --- | --- | --- | --- |
| **GDF-15** | 1.000 |  |  |  |  |  |  |  |  |  |  |  |  |  |  |
| **ACE2** | 0.304 | 1.000 |  |  |  |  |  |  |  |  |  |  |  |  |  |
| **ACE2**<sup>2-ΔΔCt</sup> | 0.395 | 0.370 | 1.000 |  |  |  |  |  |  |  |  |  |  |  |  |
| **ACE2**<sup>2-ΔΔCt#</sup> | 0.102 | 0.273 | 0.157 | 1.000 |  |  |  |  |  |  |  |  |  |  |  |
| **HIF1A**<sup>2-ΔΔCt</sup> | -0.163 | 0.021 | -0.177 | **0.704\*\*** | 1.000 |  |  |  |  |  |  |  |  |  |  |
| **HIF1A**<sup>2-ΔΔCt#</sup> | -0.262 | 0.104 | **0.637\*** | -0.062 | -0.257 | 1.000 |  |  |  |  |  |  |  |  |  |
| **AST** | 0.056 | 0.110 | -0.183 | -0.079 | 0.138 | -0.127 | 1.000 |  |  |  |  |  |  |  |  |
| **ALT** | 0.291 | 0.005 | -0.080 | -0.123 | 0.040 | 0.021 | **0.537\*** | 1.000 |  |  |  |  |  |  |  |
| **GGT** | 0.157 | 0.144 | -0.140 | -0.218 | -0.078 | -0.017 | 0.098 | **0.544\*** | 1.000 |  |  |  |  |  |  |
| **LDH** | -0.213 | -0.187 | 0.186 | 0.165 | 0.198 | -0.305 | 0.007 | -0.155 | -0.079 | 1.000 |  |  |  |  |  |
| **D-Dimer** | **0.592\*** | 0.422 | **0.748\*\*** | 0.239 | -0.212 | -0.233 | -0.064 | 0.068 | 0.123 | 0.458 | 1.000 |  |  |  |  |
| **HCT** | **0.448\*** | -0.425 | **-0.485\*** | -0.373 | -0.181 | 0.037 | 0.284 | 0.028 | -0.191 | 0.184 | -0.249 | 1.000 |  |  |  |
| **Hb** | -0.402 | -0.219 | **-0.511\*** | -0.308 | -0.033 | 0.103 | 0.223 | 0.014 | -0.199 | 0.090 | -0.343 | **0.912\*\*\*** | 1.000 |  |  |
| **RBC** | -0.344 | -0.347 | -0.397 | -0.324 | -0.118 | 0.001 | 0.356 | -0.046 | -0.130 | 0.187 | -0.202 | **0.905\*\*\*** | **0.779\*\*\*** | 1.000 |  |
| **Igg** | -0.113 | -0.202 | -0.341 | -0.469 | -0.320 | -0.289 | 0.587 | 0.377 | -0.230 | -0.026 | -0.148 | 0.457 | 0.150 | 0.525 | 1.000 |

Correlation coefficient and p-value, \*p < 0.05; \*\*p < 0.01; \*\*\*p < 0.001. 2<sup>-ΔΔCt</sup> (delta delta CT, relative fold change expression in plasma); 2<sup>-ΔΔCt#</sup> (delta delta CT, relative fold change expression in buffy coats); ACE2: angiotensin 2-converting enzyme; GDF15: growth differentiation factor 15; HIF1A: hypoxia inducible factor 1 alpha; AST: aspartate aminotransferase; ALT: alanine aminotransferase; GGT: gamma glutamyl transpeptidase; LDH: lactate dehydrogenase; HCT: hematocrit; HB: hemoglobin; RBC: Red Blood Cells; Igg: Immunoglobulin G.

**Supplemental Table 5.** ACE2 genotypes frequency among the study population.

**Supplemental Table 6.** Association between ACE2 genotypes with circulating levels of GDF15 and ACE2

